# TrkA^+^ sensory neurons regulate osteosarcoma proliferation and vascularization to promote disease progression

**DOI:** 10.1101/2024.06.20.599869

**Authors:** Qizhi Qin, Sowmya Ramesh, Zhao Li, Lingke Zhong, Masnsen Cherief, Mary Archer, Xin Xing, Neelima Thottappillil, Mario Gomez-Salazar, Mingxin Xu, Manyu Zhu, Leslie Chang, Ankit Uniyal, Khadijah Mazhar, Monisha Mittal, Edward F. McCarthy, Carol D. Morris, Benjamin Levi, Yun Guan, Thomas L. Clemens, Theodore J. Price, Aaron W. James

## Abstract

Bone pain is a presenting feature of bone cancers such as osteosarcoma (OS), relayed by skeletal-innervating peripheral afferent neurons. Potential functions of tumor-associated sensory neurons in bone cancers beyond pain sensation are unknown. To uncover neural regulatory functions, a chemical-genetic approach in mice with a knock-in allele for TrkA was used to functionally perturb sensory nerve innervation during OS growth and disease progression. TrkA inhibition in transgenic mice led to significant reductions in sarcoma-associated sensory innervation and vascularization, tumor growth and metastasis, and prolonged overall survival. Single-cell transcriptomics revealed that sarcoma denervation was associated with phenotypic alterations in both OS tumor cells and cells within the tumor microenvironment, and with reduced calcitonin gene-related peptide (CGRP) and vascular endothelial growth factor (VEGF) signaling. Multimodal and multi-omics analyses of human OS bone samples and human dorsal root ganglia neurons further implicated peripheral innervation and neurotrophin signaling in OS tumor biology. In order to curb tumor-associated axonal ingrowth, we next leveraged FDA-approved bupivacaine liposomes leading to significant reductions in sarcoma growth, vascularity, as well as alleviation of pain. In sum, TrkA-expressing peripheral neurons positively regulate key aspects of OS progression and sensory neural inhibition appears to disrupt calcitonin receptor signaling (CALCR) and VEGF signaling within the sarcoma microenvironment leading to significantly reduced tumor growth and improved survival. These data suggest that interventions to prevent pathological innervation of osteosarcoma represent a novel adjunctive therapy to improve clinical outcomes and survival.

## Main

Bone pain is a near universal feature of bone cancers such as osteosarcoma (OS)^1,2^, relayed by skeletal-innervating nociceptive neurons^3^. Beyond sensation of pain, potential efferent / regulatory roles of skeletal-innervating neurons in bone cancers are unknown. Since the visualization of peripheral neurons in cancer tissues in the late 1800s^4^, accumulating data in tumor biology suggests that tumor-associated neurons regulate epithelial-derived cancers^5^, such as breast, prostate and pancreatic carcinomas^6^. In contrast, neurosecretory functions of peripheral innervation in tumors of mesenchymal origin (sarcomas) remain unknown.

The skeleton is predominantly innervated by peripheral afferent (sensory) nerves^7,8^. Recent studies from our laboratory and others have demonstrated an essential role for peripheral nerves that innervate the skeleton in the regulation of skeletal cells and tissues^9–16^. Most skeletal neurons represent NGF (nerve growth factor) responsive TrkA (Tropomyosin receptor kinase A) expressing fibers^10^, which interact with bone tissues to influence skeletal development^13,17^ and tissue repair^9,10,14,18^. More recently, we reported that sensory nerves of the skeleton play similar important roles in pathologic processes of bone. For example, TrkA^+^ axons sprout around the benign/reactive bone tumor known as heterotopic ossification, to positively regulate the disease process^11^. These aggregate findings in non-neoplastic bone tissue strongly implicate peripheral nerves of the skeleton as niche regulators across orthopedic contexts and led us to examine the role of skeletal innervation in sarcomas of bone.

Here, transgenic animals were employed to visualize and inactivate TrkA^+^ nerves in a mouse orthotopic OS xenograft model. Our results show that aberrant sprouting of TrkA^+^ nerves accelerate key aspects of OS tumor pathophysiology, altering cellular phenotypes within the sarcoma proper and the surrounding tumor microenvironment. Moreover, targeting pathological innervation in OS using FDA-approved agents was identified as a novel and effective adjunctive therapy to improve disease progression in sarcoma patients.

## Results

### Pathological nerve sprouting is a consistent feature of osteosarcoma

In order to map the sensory nerve-tumor interactions in osteosarcoma (OS), we first utilized an orthotopic xenograft model of 143B tumor cells in immunocompromised neuronal reporter mice (*Thy1*-YFP;NOD-*Scid*) **(Fig. 1a)**. Normal innervation patterns were first confirmed, focusing on the uninoculated tibial periosteum **(Fig. 1b)**. Consistent with prior reports^7^, thin linear innervation was present in non-neoplastic periosteum as highlighted by Beta-III tubulin (TUBB3) immunostaining, as well as staining for the peptidergic marker (CGRP, Calcitonin gene-related peptide), and sympathetic marker (TH, Tyrosine hydroxylase). Next, similar analyses were performed using 143B OS implants **(Fig. 1c-i)**. Adjacent to OS implants, a high density of TUBB3^+^ and *Thy1*^+^ nerve fibers were observed **(Fig. 1c-e)**. The majority of peri-tumoral nerves showed TrkA (tropomyosin-related kinase A), CGRP, and NF200 (neurofilament 200) immunostaining, **(Fig. 1f-h)**, while the minority of nerves appeared to be immunoreactive for the sympathetic marker TH **(Fig. 1i)**. Confirming this observation, quantification of pathological peri-tumoral nerve sprouting showed higher fold increase in CGRP^+^ sensory axons as compared to TH^+^ sympathetic axons **(Fig. 1j)**. Spatial mapping of nerve-OS interaction using Imaris software was performed **(Extended Data Fig. 1a)**, which showed that most nerves resided in close proximity to the tumor leading edge in the peri-tumoral area **(Extended Data Fig. 1b, c)**. Re-analysis of the publicly available RNA sequencing EMBL-EBI database^19,20^ confirmed high expression of neurotrophins, such as nerve growth factor (*Ngf*) and brain-derived neurotrophic factor (*Bdnf*) **(Fig. 1k)**, along with other axon guidance molecules **(Extended Data Fig. 1d)** across multiple human OS cell lines. Sarcoma-derived NGF was further verified by immunostaining in xenograft OS implants, which showed a 112-fold increase in comparison to non-neoplastic tibial bone **(Fig. 1l-n)**. In summary, pathological sensory innervation associated with axon guidance cues, including NGF expression by tumor cells, represents a consistent feature in OS.

**Fig. 1.**
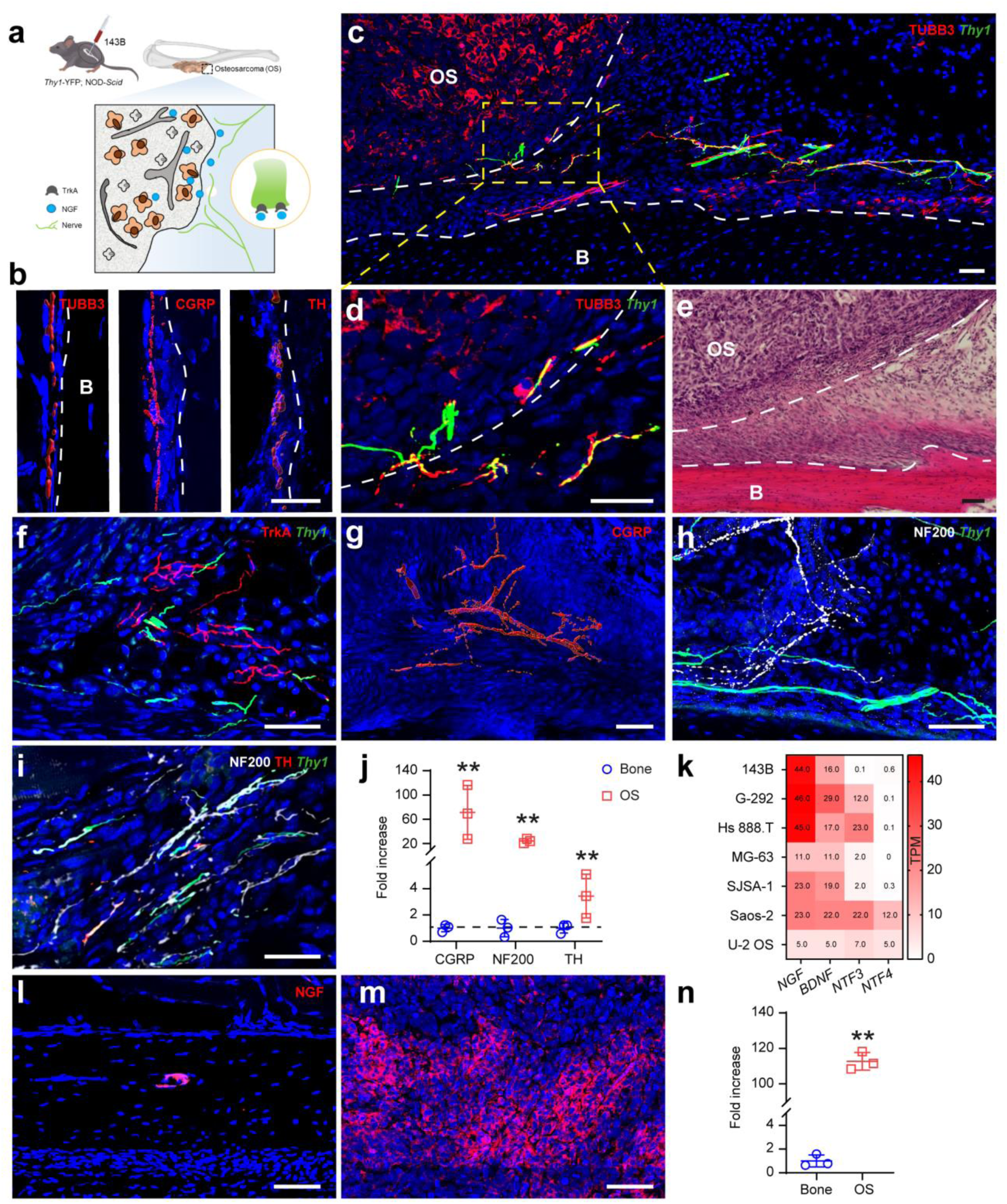
Pathological nerve sprouting in osteosarcoma (OS) and neuron-to-OS interaction. Sarcoma implants generated by orthotopic implantation of 143B osteosarcoma cells (1×10^6^ cells) in *Thy1*-YFP; NOD-*Scid* reporter mice. Tumors harvested 7 d after implantation. **a**, Schematic of the study. **b**, Representative immunofluorescence staining of Beta III tubulin (TUBB3), Calcitonin gene-related peptide (CGRP), and Tyrosine hydroxylase (TH)-expressing nerves in normal tibial periosteum. B: bone. **c**, Tile scan showing OS implant, bone, and immunofluorescence staining of pan-neuronal marker TUBB3 (red) and *Thy1*-YFP reporter activity (green). White dashed line represents the tumor boundary and the boundary between the periosteum and the underlying cortical bone. OS: osteosarcoma. **d**, High magnification image from **c**. **e**, Representative histologic appearance by routine H&E-stained section, showing a well-circumscribed OS tumor adjacent to the tibial cortex below. **f-i**, Representative immunofluorescence staining of neuron markers including **f**, Tropomyosin receptor kinase A (TrkA). **g,** CGRP. **h**, Neurofilament 200 (NF200), and **i,** TH, along with *Thy1* reporter adjacent to the sarcoma implants. **j**, Fold change in CGRP, NF200 and TH immunostaining around OS tumors in comparison to normal tibial periosteum. Dashed line indicates baseline innervation in normal periosteum. **k**, Expression levels of *Nerve growth factor (NGF), Brain-derived neurotrophic factor (BDNF), Neurotrophin 3 (NTF3)*, and *Neurotrophin 4 (NTF4)* in human OS cell lines, shown by heatmap. **l**, **m**, Immunostaining and **n**, Quantification of NGF in non-neoplastic bone and 143B OS implants. N=3 for all immunostainings. Scale bar: 50 µm. For scatterplots, data are expressed as the mean ± 1 SD. Individual dots in scatterplots represent values from single measurements. Statistical analysis was performed using an unpaired two-way Student’s t test. \*\**p* < 0.01.

### Inhibition of TrkA neuronal signaling mitigates OS growth and disease progression

Skeletal innervating sensory neurons commonly express the high-affinity NGF receptor TrkA^21^. Prior studies from our group utilized a chemical-genetic approach to temporally inhibit TrkA catalytic activity, in which TrkA^F592A^ mice are homozygous for a phenylalanine-to-alanine point mutation (F592A) in exon 12 of the mouse *Ntrk1* gene^22^ allowing for TrkA kinase inhibition by the membrane-permeable small molecule 1NMPP1, while TrkA^WT^ mice are insensitive^18,22^. TrkA^WT^ and TrkA^F592A^ mice were crossbred with NOD-*Scid* mice to allow sarcoma xenografting. Next, 143B human OS cells were implanted orthotopically in the proximal tibia^23,24^ in TrkA^WT^; NOD-*Scid* or TrkA^F592A^; NOD-*Scid* mice (hereafter denoted as TrkA^WT^ and TrkA^F592A^ mice), and TrkA inhibition was achieved by 1NMPP1 administration according to previously validated schedules^25^ **(Fig. 2a)**. First, reduction of innervation among TrkA^F592A^ mice was confirmed by TUBB3 immunostaining in the peri-tumoral area, (88.3% reduction in TUBB3 staining, **Fig. 2b)**. Next, *in vivo* tumor growth was assessed by combined approaches including bioluminescence imaging using luciferase activity, ^18^F-FDG PET-CT, and serial caliper measurements **(Fig. 2c-e, Extended Data Fig. 2a)**. In comparison to TrkA^WT^ mice, tumor growth was significantly impeded among TrkA^F592A^ mice, including a significant decrease in luciferase activity (**Fig. 2c**, 60.0% reduction), FDG-PET **(Fig. 2d**, 28.8% reduction), and tumor size **(Fig. 2e**, 51.5% reduction at study endpoint). Of note, changes in tumor growth could not be explained by a direct effect of small molecule 1NMPP1 on 143B cells, which showed no effect on *in vitro* cell viability **(Extended Data Fig. 2b)**. Histologic examination of tumor implants demonstrated decreased intra-tumoral proliferation **(Fig. 2f**, 81.0% reduction in Ki67 staining) and increased apoptosis **(Fig. 2g**, 4.2-fold increase in TUNEL labeling) among TrkA^F592A^ mice. In order to confirm the impact of sensory innervation on invasion and metastasis, pulmonary metastatic burden was assessed by human nuclear antigen (HNA) staining in serial cross-sections of lung tissues **(Fig. 2h)**. Results indicated a notable reduction of lung metastasis among TrkA^F592A^ mice at 28 d after orthotopic tumor cell inoculation. The overall pulmonary burden of disease was quantified by assessing the overall number of metastatic foci and overall burden of HNA^+^ cells per cross-section of lung tissue. A conspicuous 86.5% decrease in metastatic foci and 43.4% reduction in pulmonary HNA total cell burden was observed among TrkA^F592A^ mice **(Fig. 2i)**. Consistent with these findings and in a separate cohort of animals, mice with inhibition of TrkA^+^ neurons demonstrated prolonged overall survival **(Fig. 2j, 42.7% increase in median overall survival)**. Together, these data demonstrate that inhibition of TrkA neuronal signaling significantly slows OS tumor growth and reduces metastatic spread to enhance overall survival.

**Figure 2.**
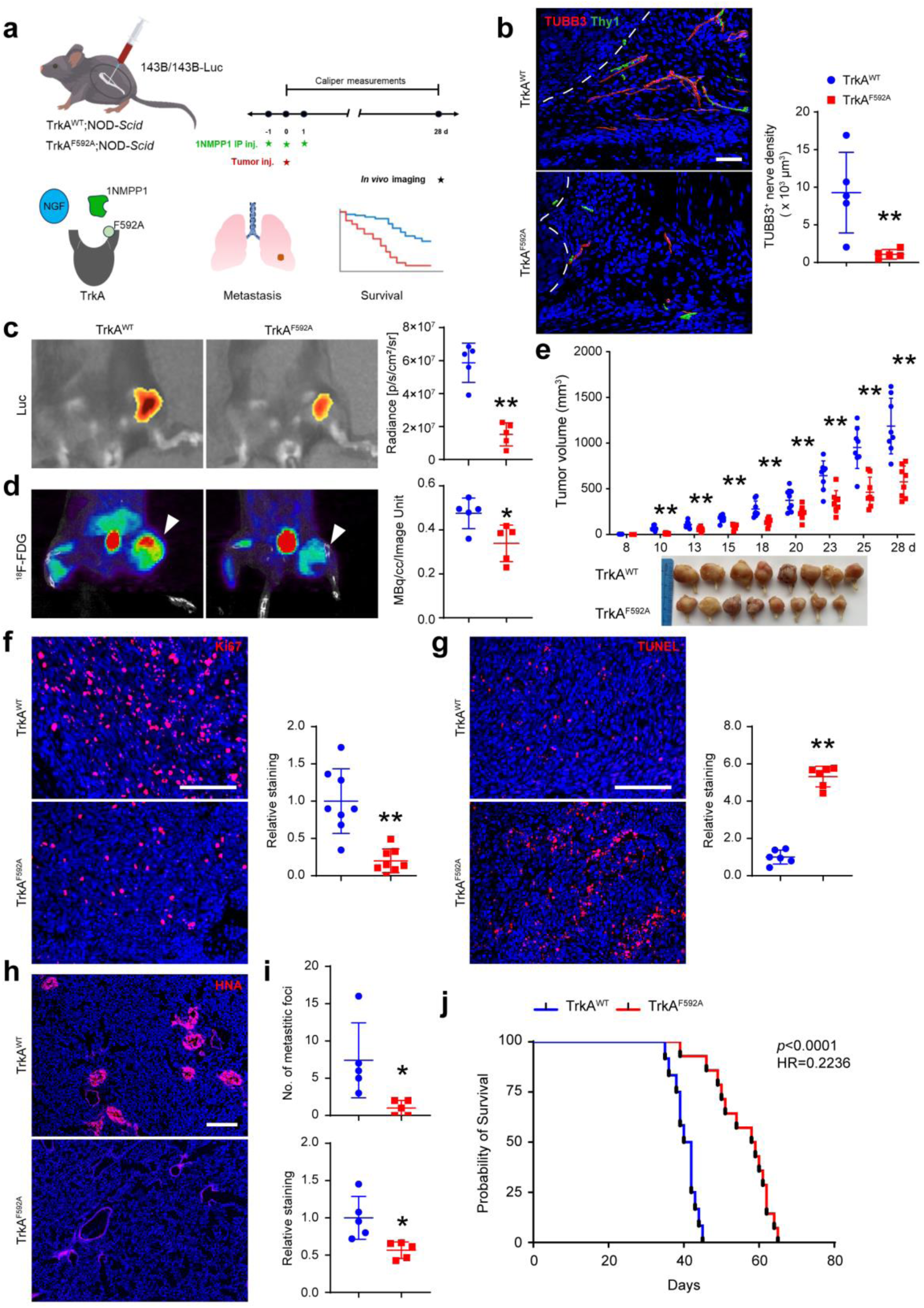
Inhibition of TrkA catalytic activity mitigates OS disease progression. **a**, Schematic of study design. TrkA inhibition performed using 1NMPP1 treatment in TrkA^F592A^;*Thy1*-YFP;NOD-*Scid* transgenic mice (termed TrkA^F592A^). Control littermates were TrkA^WT^;*Thy1*-YFP;NOD-*Scid* (TrkA^WT^), which also received 1NMPP1. **b**, Peri-tumoral innervation examined by TUBB3 immunostaining and semi-quantification. Dashed white lines indicate tumor edge. **c**, Bioluminescence imaging among TrkA^WT^ and TrkA^F592A^ mice (left) and quantification of imaging signaling intensity (right, photons/sec/cm^2^/steradian) at 28 d after 143B-Luc cell implantation. **d**, ^18^F-FDG PET-CT images (left) and quantification (right) at 28 d after 143B cell implantation. **e**, Tumor volume calculated by caliper measurements until 28 d post-implantation in TrkA^WT^ and TrkA^F592A^ mice (above) and gross pathology of all tumors (below). **f**, Proliferation assessed by Ki67 immunostaining and quantification. **g**, Apoptosis assessed by TUNEL staining and quantification. **h**, Lung metastasis assessed by Human Nuclei Antigen (HNA) immunostaining on cross-sections of pulmonary fields. **i**, Quantification of metastatic foci and semi-quantification of HNA staining. **j**, Overall survival evaluated using Kaplan–Meier curves (*N*=10 TrkA^WT^ and *N*=13 TrkA^F592A^ mice). Scale bar: 50 µm in **b**, 100 µm in **f-h**. *N*=5-6 animals per group for **b-d**, **g-i**, *N*=8 animals per group for **e**, **f.** Data are expressed as the mean ± 1 SD. Individual dots in scatterplots represent values from single animal measurements. Statistical analysis was performed using an unpaired two-way Student’s t test. \**p* < 0.05, \*\**p* < 0.01.

### TrkA neuronal inhibition alters the transcriptional landscape of osteosarcoma at the single-cell level

To understand the mechanism by which TrkA neuronal inhibition led to decreased sarcoma growth, single-cell RNA sequencing (scRNA-seq) of TrkA^WT^ and TrkA^F592A^ xenograft 143B OS implants were performed **(Fig. 3a)**. Unsupervised clustering identified seven cell clusters, including human tumor cells (expressing *TMSB10*, *HMGA1*) and six groups of mouse tumor-associated cell types constituting the tumor microenvironment (TME) **(Fig. 3b)**. The TME consisted of macrophages (expressing *Apoe*, *Cd74*), fibroblasts (expressing *Dcn, Col3a1*), neutrophils (expressing *Acod1*, *S100a8*), two clusters of lymphatic/endothelial cells (LEC, expressing *Pecam1*, *Lyve1*), myogenic cells (expressing *Acta1*, *Mylpf*), and NK cells (expressing *Gzma*, *Nkg7*) **(Fig. 3c)**. Each cluster was evenly represented across both TrkA^WT^ and TrkA^F592A^ mice **(Supplementary Table S1, Supplementary Fig. S1a)**. Gene transcripts of *NGF* among tumor cells **(Supplementary Fig. S1b)** and TME cells **(Supplementary Fig. S1c)** remained unchanged by TrkA inhibition. The downstream consequences in response to TrkA inhibition were sequentially assayed by examining tumor cells and TME cells separately using unbiased GO term analysis. Among tumor cells and TME cells from TrkA^WT^ implants, network clustering of the enriched genes applying the Markov Clustering Algorithm (MCL) revealed eight **(Fig. 3d, Supplementary Table S2)** and three clusters **(Fig. 3e, Supplementary Table S3)** of functionally related terms, respectively. The results of GO analysis of differentially expressed genes showed that the products of these genes were predominantly associated with axon extension and angiogenesis among tumors cells and TME cells harvested from TrkA^WT^ implants. GO analysis of tumor cells and TME cells from TrkA^F592A^ implants resulted in seven **(Supplementary Fig. S2a, Supplementary Table S4)** and five clusters, respectively **(Supplementary Fig. S2b, Supplementary Table S5),** with divergent terms from Semaphorin Plexin signaling to BMP signaling. Thus, GO term analysis of xenograft tumor implants identified broad-scale derangements in tumor and TME transcriptome after TrkA inhibition, with many terms linking to the neuro-vasculature.

**Figure 3.**
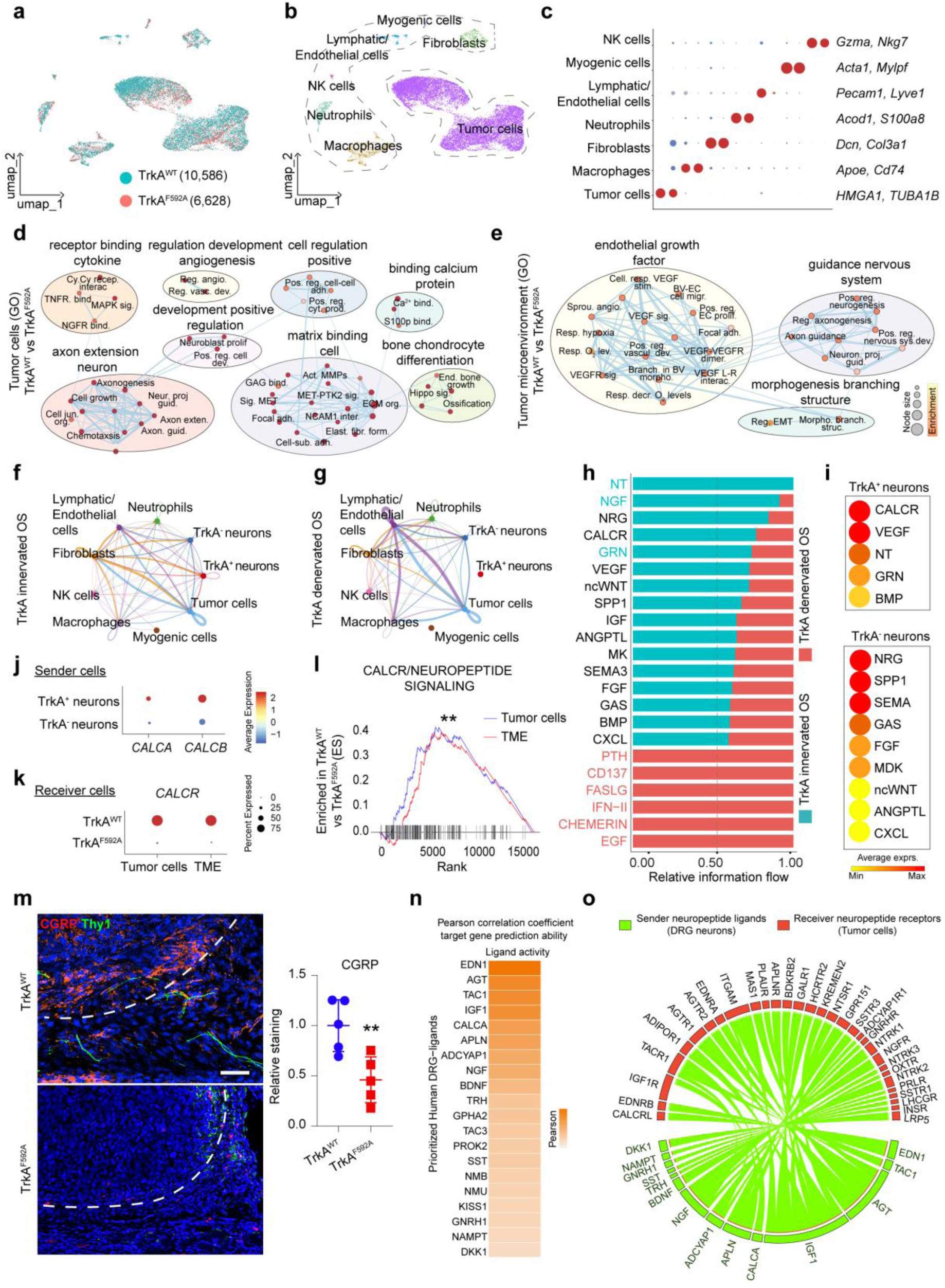
Single-cell transcriptomics identifies altered cellular processes and impaired CALCR/neuropeptide signaling with TrkA inhibition. Single cell RNA sequencing of the tumor site performed 12 d after implantation of 143B OS cells into either TrkA^F592A^;*Thy1*-YFP;NOD-*Scid* (TrkA^F592A^) or TrkA^WT^;*Thy1*-YFP;NOD-*Scid* (TrkA^WT^) mice. All animals received small molecule 1NMPP1. **a,** Reduced-dimensionality (UMAP) visualization and clustering of xenograft tumor implants harvested from TrkA^WT^ (10,586 cells) and TrkA^F592A^ mice (6,628 cells), pooled from three mice per genotype. **b,** UMAP projection and clustering identifying cell clusters, including human tumor cells (13,170 cells), and mouse tumor microenvironment cells (TME, 3,837 cells). **c,** Dot plots of characteristic gene markers among each cell cluster. Network of enriched GO terms and Reactome pathways generated with g:Profiler and EnrichmentMap using Cytoscape among **d,** tumor cells implanted in TrkA^WT^ mice, and **e,** TME cells from TrkA^WT^ mice. Each node (circle) represents a gene set characterized by a particular GO term or reactome pathway. The node fill indicates the enrichment score (FDR q-value). The thickness of blue lines (edges) indicates the number of shared genes (overlap) between two connected nodes. Nodes with high overlap are clustered together, forming groups characterized by similar terms and pathways. Networks of cell-cell inferred interactions show the number of ligand-receptor pairs (edges) between each cell type (nodes) among either **f,** TrkA innervated OS or **g**, TrkA denervated OS implants. The width of the edge indicates the number of ligand-receptor interactions between cell types**. h,** Overall relative information flow by signaling pathway among TrkA innervated OS versus TrkA denervated OS. Significantly enriched pathways are colorized. **i,** Circle heatmaps showing predominantly enriched signaling pathways within TrkA^+^ and TrkA^-^ neurons^26^. **j,** Expression of corresponding *CALCA* and *CALCB* ligands among the DRG neuronal sender cells (TrkA^+^ and TrkA^-^ neurons)^26^. k, *CALCR* receptor gene expression among the receiver cells (tumor cells and TME cells) under TrkA^WT^ versus TrkA^F592A^ conditions. **l,** Gene set enrichment analysis of CALCR/neuropeptide signaling pathway among tumor cells and TME cells under TrkA^WT^ versus TrkA^F592A^ conditions. **m,** CGRP immunostaining, along with *Thy1*-YFP reporter activity adjacent to the sarcoma implants among TrkA^WT^ and TrkA^F592A^ OS implants and corresponding quantification. Dashed white lines indicate tumor edge. **n,** Heatmap showing the ligand activity of top-predicted neural derived ligands^28^ ranked according to Pearson correlation. **o,** Circos plot summarizing the NicheNet interactome analysis that identified potential signaling connections between neural derived ligands (neuropeptides + neurotrophins) expressed by (sender) human DRG neurons^28^ (n=3,927 cells) and the paired target receptors expressed by (receiver) tumor cells (n=8,344 cells) harvested from TrkA^WT^ OS implants. For all graphs, data are expressed as the mean ± 1 SD. \**p* < 0.05, \*\**p* < 0.01.

To computationally reconstruct the paracrine cellular communication more precisely between TrkA^+^ neurons and tumor cells, available single-nuclei RNA-sequencing (sNuc-seq) from human lumbar dorsal root ganglion (DRG) neurons was utilized which would naturally innervate the skeleton of the lower extremity^26^. The computationally merged dataset comprised of DRG resident neurons expressing *NTRK1*^26^ **(Supplementary Fig. S3a, b)**, tumor cells and TME cells, referred to as ‘TrkA innervated OS’ **(Extended Data Fig. 3a, b)**. Similarly, another dataset was computationally integrated by combining only TrkA^-^ neurons, tumor cells, and TME cells, referred to as ‘TrkA denervated OS’**(Extended Data Fig. 3c, d)**. These TrkA^+^ neurons largely represent nociceptive neurons with high expression of *TAC1* **(Supplementary Fig. S3c)** and *CALCA* **(Supplementary Fig. S3d)**.

The R package CellChat^27^ was used to investigate the overall cell-cell network interactions among different types of TME cells, tumor cells with TrkA^+^ neurons (**Fig. 3f)** and TrkA^-^ neurons **(Fig. 3g)**. Using relative information flow, overall differences in cellular communication between the two datasets were inferred **(Fig. 3h)**. Among the TrkA innervated OS, the top six pathways that were enriched in their communication were neurotrophin (NT pathway), NGF pathway, neuregulin (NRG pathway), calcitonin receptor signaling (CALCR pathway), granulin pathway, and vascular endothelial growth factor signaling (VEGF pathway) **(Fig. 3h)**. Upon dissecting the signaling pathways using module scores that were predicted for enrichment within TrkA^+^ neurons as sender cells, CALCR, VEGF, and NT signaling pathways were among the top three **(Fig. 3i)**. In contrast, other signaling pathways such as NRG signaling, secreted phosphoprotein-1 (SPP1) signaling, and semaphorin signaling were all more predicted to be derived by TrkA^-^ rather than TrkA^+^ neurons as sender cells **(Fig. 3i)**. As expected, gene transcripts for CGRP ligands (*CALCA*, *CALCB*) were enriched among TrkA^+^ neurons compared to non-TrkA expressing neurons **(Fig. 3j)**. Likewise, module scores of curated gene sets confirmed a decrease in transcripts associated with CALCR signaling activation among TrkA^F592A^ implants, both when examining tumor cells as well as TME cells **(Fig. 3k)**; (key individual genes shown in **Supplementary Fig. S4)**. Having identified candidate signaling pathways predominantly facilitated by TrkA^+^ neurons, gene set enrichment analysis for CALCR/neuropeptide signaling was performed among TrkA^WT^ or TrkA^F592A^ OS single-cell datasets **(Fig. 3l).** Results showed an overall significant enrichment score (p<0.001) for CALCR/neuropeptide signaling among cells derived from TrkA^WT^ as compared to TrkA^F592A^ mice, both when examining tumor cells as well as TME cells **(Fig. 3l).**

Next, these computational findings were further substantiated using our additional sequencing datasets of human nociceptive neurons, as assayed using sNuc-seq^28^ and spatial transcriptomics.^29^ Here, 12 cell clusters of human DRG neurons^28^ collected from lumbar L1-L5 levels **(Extended Data Fig. 4a)** were evaluated for *NTRK1* gene expression **(Extended Data Fig. 4b)**, both by sNuc-seq (average fold expression: 29.91) and spatial sequencing (average fold expression: 37.17, **Extended Data Fig. 4c, c’)**. Across both sNuc-seq and spatial sequencing, *NTRK1* gene transcripts were differentially expressed among various neuronal subclusters; PENK+ nociceptors (average fold expression: 2.36), Putative silent nociceptors (average fold expression: 0.39), Aβ-nociceptors (average fold expression: 2.40), and Aδ-HTMRs (average fold expression: 6.55) (See corresponding **Source File** for the full list). Next, relative information flow was again performed, now by integrating independently generated human sNuc-seq dataset with increased depth of coverage^28^ (stratified by *NTRK1* expression) and TrkA xenograft OS dataset **(Extended Data Fig. 4d)**. Interaction analysis reiterated substantial enrichment of CALCR signaling pathway, VEGF signaling pathway, and NT signaling pathway among the TrkA^WT^ OS implants in the presence of TrkA^+^ positive neurons **(Extended Data Fig. 4d)**. Next, alterations in CALCR signaling pathway across multiple transcriptomics datasets were validated by returning to tissue sections of xenograft OS implants with and without TrkA inhibition. Immunostaining for CGRP showed significant reductions among TrkA^F592A^ animals in comparison to TrkA^WT^ mice, found in adjacency to *Thy1* reporter activity at the tumor periphery **(Fig. 3m**, 54.0% reduction). To further deduce the potential neural derived ligands that may drive the changes in CALCR/neuropeptide signaling, interactome analysis was performed. NicheNet^30^ analysis predicted a range of putative neuropeptides/ neurotrophins ligands from human DRG neurons^28^ **(Fig. 3n)**. Of these ligands, *TAC1*, *CALCA*, *NGF*, and *BDNF* were predicted to act on tumor cells, in a paracrine fashion **(Fig. 3o, Extended Data Fig. 4e**. See corresponding **Source File** for the full list). Across both sNuc-seq^28^ **(Extended Data Fig. 4f)** and spatial sequencing datasets^29^ **(Extended Data Fig. 4g)** enrichment of *CALCA* across various neuronal subclusters were verified. *CALCA* was enriched among putative silent nociceptors (average fold expression: 20.30), Aδ-HTRMs (average fold expression: 13.76), PENK+ nociceptors (average fold expression 11.24), and Aβ-nociceptors (average fold expression: 5.03, **Extended Data Fig. 4e**). Thus, impaired innervation in the context of the TrkA neuronal signaling inhibition was associated with downstream reductions in CGRP signaling across different ‘omics modalities. Further, our results implicate neuronal derived CGRP among other signaling molecules that positively regulate neuron-to-sarcoma interactions in OS disease progression.

### TrkA neuronal inhibition alters osteosarcoma associated vasculature

Nerves and vessels influence the growth and trajectory of one another during development and disease^31^. Indeed, this phenomenon was confirmed in our NOD-*Scid* xenograft OS mice model wherein a spatial relationship between Endomucin (EMCN^+^) blood vessels and NF200^+^ axons were confirmed by dual immunostaining **(Extended Data Fig. 5a)** and quantification **(Extended Data Fig. 5b)**. Upon TrkA neuronal inhibition, marked reductions in both NF200^+^ nerve fibers and EMCN^+^ vessels were found within and around TrkA^F592A^ OS implants in comparison to TrkA^WT^ tumors **(Extended Data Fig. 5c)**. To gain additional transcriptomic insights into this neurovascular disruption in response to TrkA inhibition, tumor-associated lymphatic endothelial cells (LECs) among TrkA^WT^ and TrkA^F592A^ tumor implants were re-clustered and analyzed **(Fig. 4a)**. Upon reclustering of LECs within scRNA-Seq data, two clusters of endothelial cells (Endo1, Endo2) and one cluster of lymphatic cells were identified **(Fig. 4b)**. Each cell type was annotated by known marker genes such as *Pecam1*, *Emcn, Cdh5* for Endo1; *Kdr*, *Cd34*, *Kit* for Endo2 (endothelial progenitor cluster), and *Lyve1, Pdpn, Prox1* for lymphatic vessels **(Fig. 4c)**. Each cluster was represented across both TrkA^WT^ and TrkA^F592A^ tumor implants **(Fig. 4d).** Gene module scores revealed a more proliferative **(Extended Data Fig. 5d)**, more angiogenic **(Extended Data Fig. 5e)** and a stalk-like phenotype for Endo1 compared to Endo2 **(Extended Data Fig. 5f, g)**. Pseudobulk sequencing analysis of LECs from TrkA^WT^ and TrkA^F592A^ implants demonstrated ∼200 DEGs **(Supplementary Table S6)** and heatmap analysis of the top-ranking DEGs revealed distinct transcriptomic signatures among TrkA^WT^ and TrkA^F592A^ LECs (**Fig. 4e**). Notably, CALCR signaling related genes such as Receptor activity modifying protein, *Ramp3* (log fold change 3.81, *p*<0.01), gene markers associated with endothelial tip-cell phenotype such as potassium inwardly rectifying channel subfamily J member 8 *Kcnj8* (log fold change 2.77, *p*<0.05), and Endomucin *Emcn* (log fold change 0.77, *p*<0.01) were significantly enriched among LECs derived from TrkA^WT^ in comparison to TrkA^F592A^ mice. Next, unbiased GO term enrichment analysis among TrkA^WT^ versus TrkA^F592A^ LECs resulted in three clusters of functionally related terms, including positive regulation of chemotaxis and blood vessel morphogenesis development **(Fig. 4f, Supplementary Table S7)**. GO term analysis among TrkA^F592A^ LECs is shown in **Extended Data Fig. 5h, Supplementary Table S8**. Having demonstrated global transcriptional alterations among tumor LECs in response to TrkA inhibition, we focused on neurovascular perturbations. Module scores of curated gene sets confirmed reductions in gene transcripts associated with neurotrophin signaling **(Extended Data Fig. 5i)** and CALCR signaling pathway **(Extended Data Fig. 5j)** among TrkA^F592A^ LEC compared to TrkA^WT^ LEC. Neuron-to-vascular communication was further corroborated by downstream signaling changes with reductions in angiogenesis **(Fig. 4g)** and VEGF signaling pathway **(Fig. 4h)**, among TrkA^F592A^ LECs compared to TrkA^WT^. To further investigate the degree of change in tumor-associated vasculature in response to inhibition of TrkA neuronal signaling, immunofluorescent staining for EMCN and CD31 was performed (**Fig. 4i**). Changes in innervation upon TrkA inhibition resulted in a significant disruption in tumor site vascularity among TrkA^F592A^ OS implants sites (**Fig. 4i, j,** 47.2% reduction in EMCN and 60.8% reduction in CD31 in comparison to TrkA^WT^ OS implants). Furthermore, vascular morphometric analysis showed significant reductions in total vessel length, vessel junctions, branching index and lacunarity **(Extended Data Fig. 5k-n)** among TrkA^F592A^ OS implants. Next, NicheNet^30^ was used to reconstruct potential neural-to-endothelial cell signaling by ligand-receptor matching **(Fig. 4l)**. Previously identified DRG neuronal derived pro-vasculogenic^25^ ligands (*VEGFA*, *PDGFB*, *FGF1*, *PGF*, *PDGFA*, *WNT5A*, *PDGFC*) were predicted to have interactions with neuropeptide/vascular receptors targets among tumor-associated LECs **(Fig. 4l, Extended Data Fig. 5o)**. The expression of various neuronal derived pro-vasculogenic neural derived ligands was assayed across different subclusters of human DRG neurons^28^ **(Extended Data Fig. 5p**). Of these, VEGFA ligand was highly prioritized to play a role in neural-to-endothelial paracrine interaction based on ligand activity **(Fig. 4m)**. *VEGFA* gene transcripts were enriched among putative silent nociceptors (average fold expression: 0.57). The expression of *VEGFA* (average expression: 0.19) was further verified using our spatial sequencing dataset^28^ **(Extended Data Fig. 5p)**. Lastly, these neurovascular alterations in VEGF signaling across transcriptomic datasets were substantiated by immunostaining for VEGFA among *Thy1*-YFP tumor implants with and without TrkA inhibition **(Fig. 4n)**. Tumor implants from TrkA^WT^ mice exhibited high immunoreactivity for VEGFA around the peritumoral area that coincided with *Thy1*-YFP reporter activity; while a significant reduction of VEGFA immunoreactivity was quantified among TrkA^F592A^ animals (82.6% reduction; **Fig. 4n)**. Thus, a combination of immunohistochemical methods and computational analysis identified defects in neurovascular ingrowth and implicated impaired neuron-to-vessel VEGF signaling in the context of inhibition of TrkA neuronal signaling in OS.

**Figure 4.**
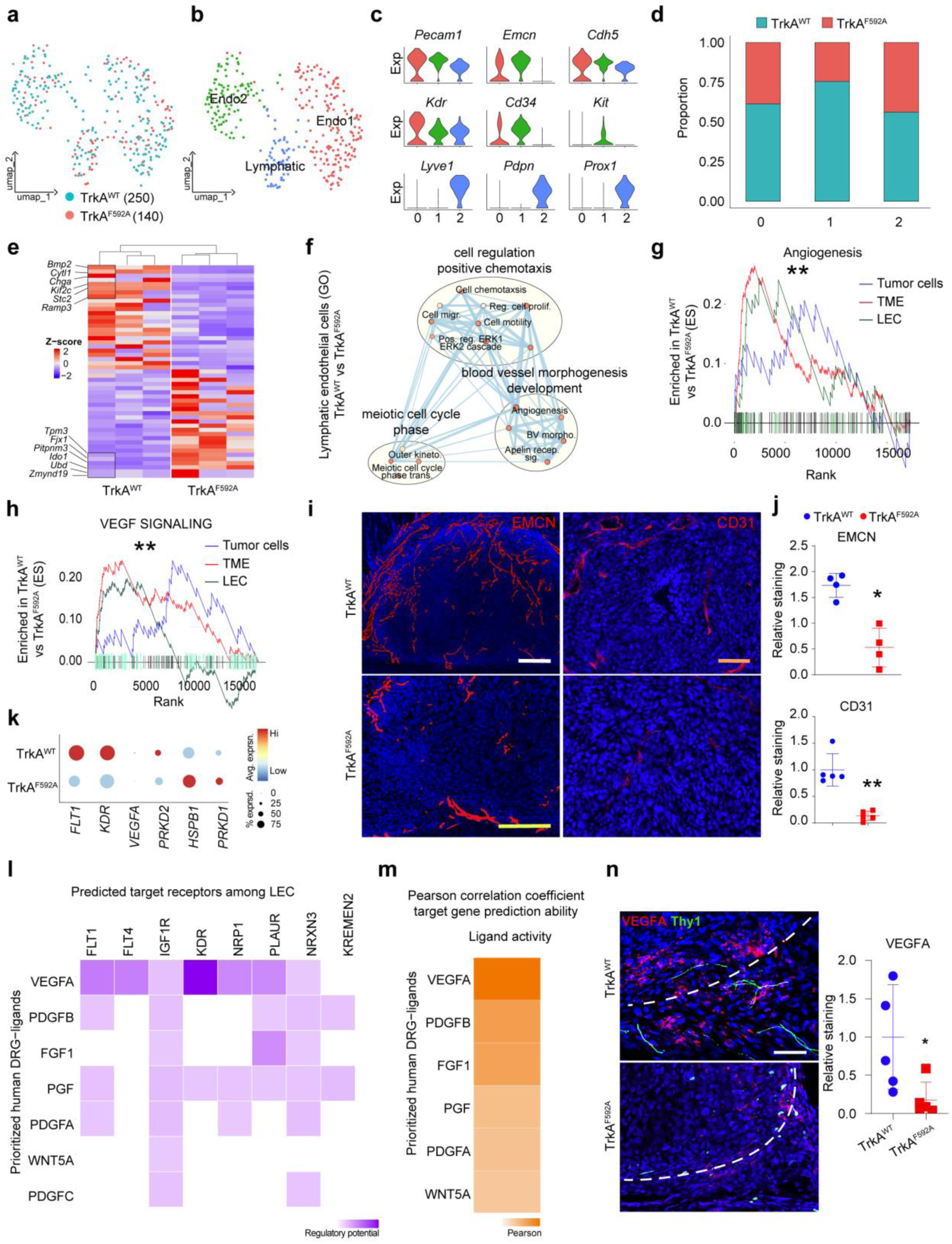
Dysregulation of tumor vascularity and VEGF signaling with deficient neural ingrowth into tumor implants. **a-h,** Single cell RNA sequencing analysis of lymphatic/endothelial cells (LEC, n=390 cells) subset from TrkA^WT^ and TrkA^F592A^ OS implants. **a,** UMAP visualization of cell clusters from TrkA^WT^ and TrkA^F592A^ mice. **b**, UMAP visualization of LEC subclusters, including two clusters of endothelial cells (endo 1, endo 2) and one cluster of lymphatic cells. **c,** Violin plots of characteristic gene markers identifying the three LEC clusters. **d,** Normalized batch distribution across subclusters and genotype. **e,** Heatmap of top DEGs among TrkA^WT^ and TrkA^F592A^ LEC. **f,** Cytoscape enriched GO terms among LECs from TrkA^WT^ vs TrkA^F592A^ mice using gProfiler. Gene set enrichment analysis score showing **g,** hallmarks of angiogenesis and **h,** reactome signaling by VEGF among tumor cells, TME cells and LECs enriched in TrkA^WT^ vs TrkA^F592A^. **i,** Immunostaining of EMCN and CD31 vessel density at d10 and d28, respectively among TrkA^WT^ and TrkA^F592A^ implants along with **j,** quantification. **k,** Dotplot showing individual genes associated with VEGF signaling among TrkA^WT^ and TrkA^F592A^ LECs. **l,** Ligand-receptor interaction plot using NicheNet analysis depicting the predicted (sender) human DRG-derived ligands^28^ with receptor target genes expressed among (receiver) TrkA^WT^ LECs. **m,** Heatmap of various human DRG neuronal^28^ derived pro-vasculogenic ligands activity ranked according to Pearson correlation coefficient. **n,** VEGFA immunostaining, along with *Thy1*-YFP reporter activity adjacent to the sarcoma implants among TrkA^WT^ and TrkA^F592A^ OS implants along with corresponding quantification. Dashed white lines indicate tumor edge. White scale bar: 50 µm; yellow scale bar: 200 µm; orange scale bar: 100 µm. N=4 animals per group for EMCN and N= 5 animals per group for CD31 immunostaining. For scatterplots, data are expressed as the mean ± 1 SD. Individual dots in scatterplots represent values from single animal measurements. Statistical analysis was performed using an unpaired two-way Student’s t test. \**p* < 0.05, \*\**p* < 0.01.

Tumor-associated macrophages (TAMs) play a vital role in altering the tumor matrix, inflammation, and vascularization^32^. However, alterations in TAMs in response to neuronal inhibition is not well-understood. To this end, TAMs among TrkA^WT^ and TrkA^F592A^ animals were analyzed by single-cell transcriptomics (**Extended Data Fig. 6a**). GO term enrichment analysis among TrkA^WT^ macrophages showed positive regulation of terms related to neuropeptide signaling, tyrosine kinase signaling and sprouting angiogenesis (**Extended Data Fig. 6b**). In contrast, TrkA^F592A^ macrophages analysis revealed enrichment of terms related to autophagy and apoptosis among other terms (**Extended Data Fig. 6b**). The frequency of peri-tumoral macrophages was examined both by sequencing (**Extended Data Fig. 6c**) as well as co-immunostaining for F4/80, a marker of murine macrophages in combination with the neuronal marker TUBB3 (**Extended Data 6d**). Consistent with the sequencing data, a significant reduction in overall peri-tumoral macrophage infiltration was observed (68.0% reduction, **Extended Data 6d**) among TrkA^F592A^ OS implants. Despite changes in macrophage frequency macrophage polarization showed no significant differences across experimental conditions using MacSpectrum **(Extended Data Fig. 6e)**. Together, these data suggest that inhibition of TrkA neuronal activity in response to OS growth is suggestive of reduction in frequency of macrophages, which may play secondary roles in regulating disease progression.

### NGF-TrkA signaling and associated peripheral innervation are conserved features among human osteosarcoma

The human relevance of NGF-TrkA signaling and OS disease progression was next examined. As a first step, NGF expression was assayed across representative human tissue sections of excised high-grade OS **(Fig. 5a)**. A previous study explored the generic alterations in the transcriptome of paired neoplastic and non-neoplastic bone in patients with OS^33^. Re-analysis of this total RNA sequencing dataset demonstrated significant enrichment for *NGF* transcripts among OS relative to the corresponding non-neoplastic bone samples (*p* = 0.007, **Fig, 5b**). Apart from *NGF*, other related axon guidance cues (such as semaphorins and other neurotrophins) were also enriched among neoplastic OS **(Supplementary Fig S5)**. Next, biological processes **(Fig. 5c, Supplementary Table S9)** and Kyoto Encyclopedia of Genes and Genomes (KEGG) pathway **(Fig. 5d, Supplementary Table S10)** analysis of upregulated genes among neoplastic OS samples demonstrated terms related to angiogenesis, axonogenesis and VEGF signaling pathway. Next, leveraging a publicly available sc-RNAseq dataset^34^ of human primary OS from six patients with various cellular composition of OS **(Fig. 5e)**, gene transcripts for *NGF* and *NTRK1* were assayed. At the single-cell level, *NGF* expression was mostly confined among osteoblastic OS cell cluster and cancer associated fibroblast clusters, compared to *NTRK1* gene transcripts, which, as expected, were lowly expressed **(Fig. 5f)**. Subclustering of all osteoblastic OS cells by patient further confirmed the expression of *NGF* across all six OS patients with variable expression of *NTRK1* **(Fig. 5g, h)**. Concordantly, immunostaining for the pan-neuronal marker TUBB3 helped to highlight the proximity of the TUBB3^+^ fine nerve fibers adjacent to human OS tumor cells, marked by RUNX2 **(Fig. 5i)**. Co-immunostaining for CD31 and RUNX2 likewise demonstrated the proximity of blood vessels around human OS tumor cells **(Fig. 5j).** Thus, human neoplastic bone phenocopies key elements of experimental findings, including expression of *NGF* among tumor cells inciting pathological axonal ingrowth among human OS-afflicted bone tissues.

**Figure 5.**
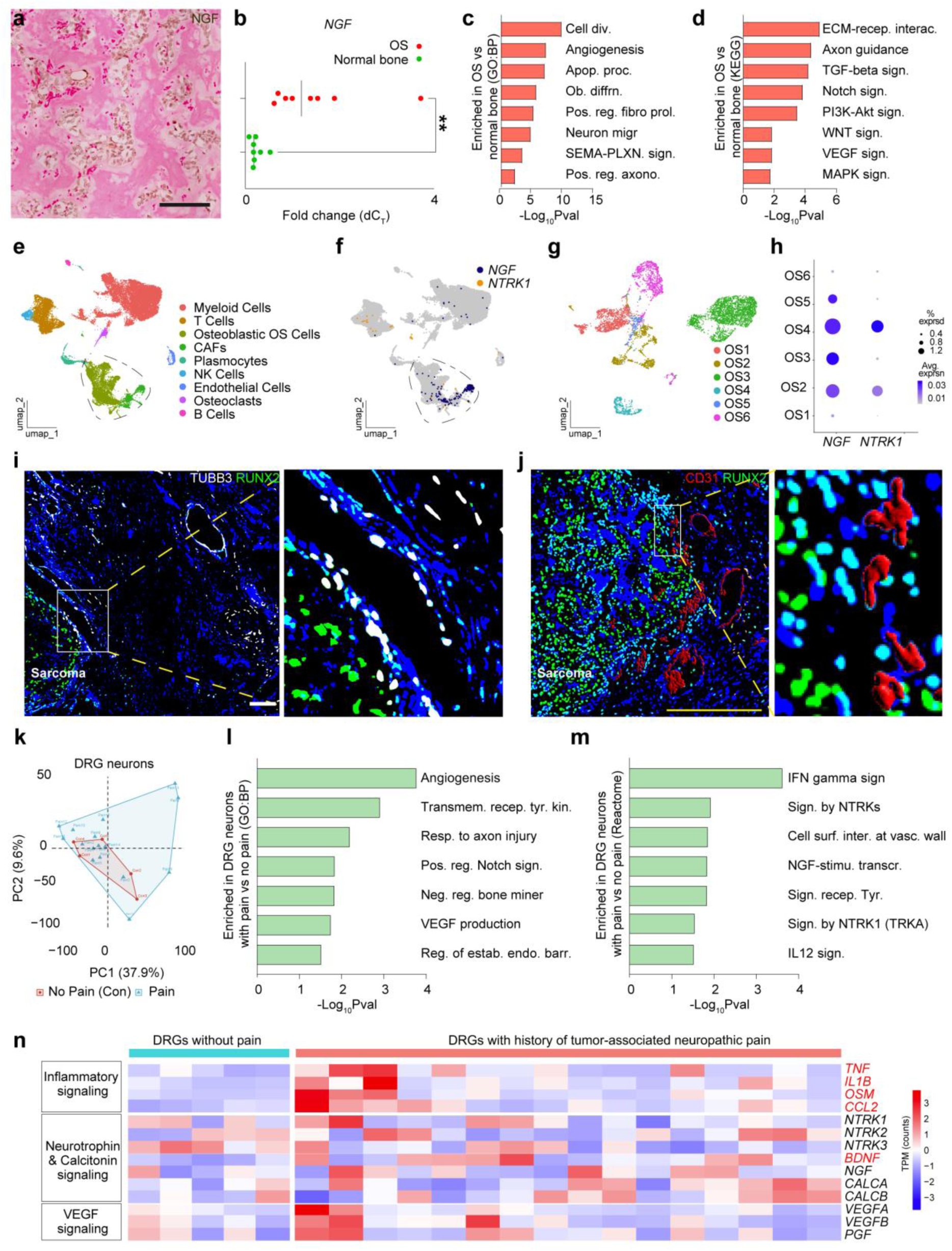
NGF-TrkA signaling and innervation within human osteosarcoma (OS) biology. **a,** Representative photomicrograph of high grade conventional human OS (N=5) immunohistochemically stained for NGF (brown). **b,** *NGF* gene expression within human OS^33^ compared to adjacent normal bone (N=8 human OS patients, *p* = 0.007). **c,** Selective GO biological process terms enriched in human neoplastic OS compared to normal bone. **d,** Selective KEGG pathways enriched in human neoplastic OS compared to normal bone. **e,** UMAP plot of six specimens of primary human OS tumors^34^. **f,** UMAP plot showing different cell types among primary human OS cells^34^. **g,** Feature plot of *NGF* and *NTRK1* gene transcripts among the primary OS cell clusters^34^. **h**, Dot plot showing expression of *NGF* and *NTRK1* gene transcripts across six human patients with primary OS^34^. **i,** Axonal growth and **j,** CD31 immunostaining within clinical samples of high grade conventional OS. Immunohistochemical staining of **i,** TUBB3^+^ nerves (white) **j**, CD31^+^ blood vessels (red), co-stained with the osteogenic transcription factor RUNX2 marking the sarcoma cells (green). **k,** Principal component analysis of human DRG neurons with and without a clinical history of tumor-associated neuropathic pain^38^. **l,** GO term analysis of enriched biological process among human DRG neurons with pain compared to those without neuropathic pain^38^. **m,** KEGG pathway analysis of terms enriched in human DRG neurons with pain compared to those without neuropathic pain^38^. **n,** Heatmap showing gene expression profile of putative inflammatory signaling, neurotrophin signaling, CALCR signaling, and VEGF signaling secretory ligands enriched among human DRG neurons associated with tumor-associated neuropathic pain^38^ (N=16 DRGs) compared to those without pain (N=5 DRGs). Ligands that are significantly enriched in DRGs neurons with neuropathic-like cancer pain are labelled in red text. \*\**p* < 0.01.

Bone cancer pain involves both a nociceptive and a neuropathic component^35^. Patients with OS have reported pain as the initial presenting symptom^36^. About 70% of the patients with OS have reported local pain^37^. After having demonstrated the spatial relationship of nerve fibers in *NGF*-expressing OS bone tissues, leveraging our bulk-RNA sequencing dataset^38^, we queried the transcriptional changes that occurred in the human DRG neurons obtained from different cancer patients with and without underlying tumor-associated neuropathic pain. Twenty-two patient DRG neuron samples were examined, of which 17 demonstrated a clinical history of neuropathic-like pain. Principal component analysis showed transcriptional variation among the sampled human DRG neurons **(Fig. 5k)**. GO term analysis using DEGs among DRG neurons associated with pain showed enriched terms in response to axon injury, and increased VEGF production **(Fig. 5l, Supplementary Table S11)**. Reactome pathway analysis further confirmed terms related to NGF transcripts and signaling by NTRK1 in those DRGs associated with pain **(Fig. 5m, Supplementary Table S12)**. This directed us to assay for putative secretory neural derived ligands of various categories including CALCR/neurotrophin signaling pathway, VEGF signaling pathway, and inflammatory signaling pathway which were indeed found to be enriched in some of the DRGs associated with history of tumor-associated neuropathic pain **(Fig. 5n)**. Notably, the neurotrophic factor *BDNF* (2.10 fold increase, *p* = 0.001) was one among the top 10 enriched neural derived ligands in those DRGs with tumor-associated neuropathic pain.

Neuropeptides including *CALCA* and pro-vasculogenic ligands such as *VEGFA, VEGFB,* and *PGF* showed increase in some of the DRG neurons with tumor-associated neuropathic pain **(Fig. 5n)**. In addition, as we previously described^38^, inflammatory markers such as *CCL2* (2.45 fold increase, *p* = 0.007), *TNF* (2.48 fold increase, *p* = 0.03), *IL1B* (4.2 fold increase, p = 0.02), and *OSM* (7.6 fold increase, *p* = 0.01) showed significant changes in those DRGs associated with pain. In sum, assessments of painful DRG neurons along with human primary OS samples is in concordance with our experimental findings, including neurotrophin/neuropeptide signaling associated with pathologic neurovascular ingrowth and neuropathic-like cancer pain.

### Bupivacaine liposomes target sensory innervation and alleviate osteosarcoma disease progression

Having observed that genetic approaches to inhibiting TrkA neuronal signaling impeded tumor growth, we sought to repurpose bupivacaine, an FDA approved non-opioid analgesic to perturb local sarcoma-associated axonal ingrowth^39^. Multivesicular liposome encapsulated bupivacaine increases the duration of local anesthetic action by slow release, available under the trade name Exparel®, has been used in >12M clinical patients (https://www.exparel.com, referred to here as L-Bup). First, *in vitro* viability assay confirmed that L-Bup had no direct cytotoxic effect on 143B tumor cells **(Fig. 6a)**. In contrast, L-Bup treatment led to significant reductions in axon length in cultured DRG neurons, consistent with the anti-axonal growth effects of Bupivacaine^40^ **(Fig. 6b)**. Next, L-Bup was administered peri-tumorally in 143B bearing NOD-*Scid* mice. As expected, L-Bup led to a significant analgesic effect on the tumor bearing limb, as measured by a decrease in paw-withdrawal frequency by von Frey filaments testing (30.7% reduction, **Fig. 6c)**, suggesting an alleviation of mechanical pain hypersensitivity. Next, L-Bup treated implants showed a significant reduction in TUBB3^+^ nerve fibers at the tumor periphery (88.1% reduction, **Fig. 6d)**. Consistently, a decrease in PGP9.5^+^ sensory nerve fiber was observed among L-Bup treated implants (82.4% reduction, **Fig. 6e)**. Strikingly, L-Bup treatment significantly reduced 143B OS tumor growth versus vehicle control by serial caliper measurements (53.6% reduction at d 28, **Fig. 6f)**. Immunostaining for Ki67 demonstrated a reduction in the intra-tumoral proliferative index of L-Bup treated implants (43.5% reduction **Fig. 6g)**. Tumor-associated angiogenesis also showed a significant decrease among L-Bup treated implants compared to control (90.6% reduction in EMCN staining, **Fig. 6h)**. Immunostaining for CGRP and VEGFA demonstrated a reduction in the secretable pool of CGRP and VEGFA among L-Bup treated implants (79.9% and 73.7% reduction, respectively, **Fig. 6i, j)**. Thus, an FDA-approved analgesic with known anti-growth effects on peripheral nerve axons leads not only to improved metrics of pain sensation, but also attenuates sarcoma innervation, and impedes tumor progression.

**Figure 6.**
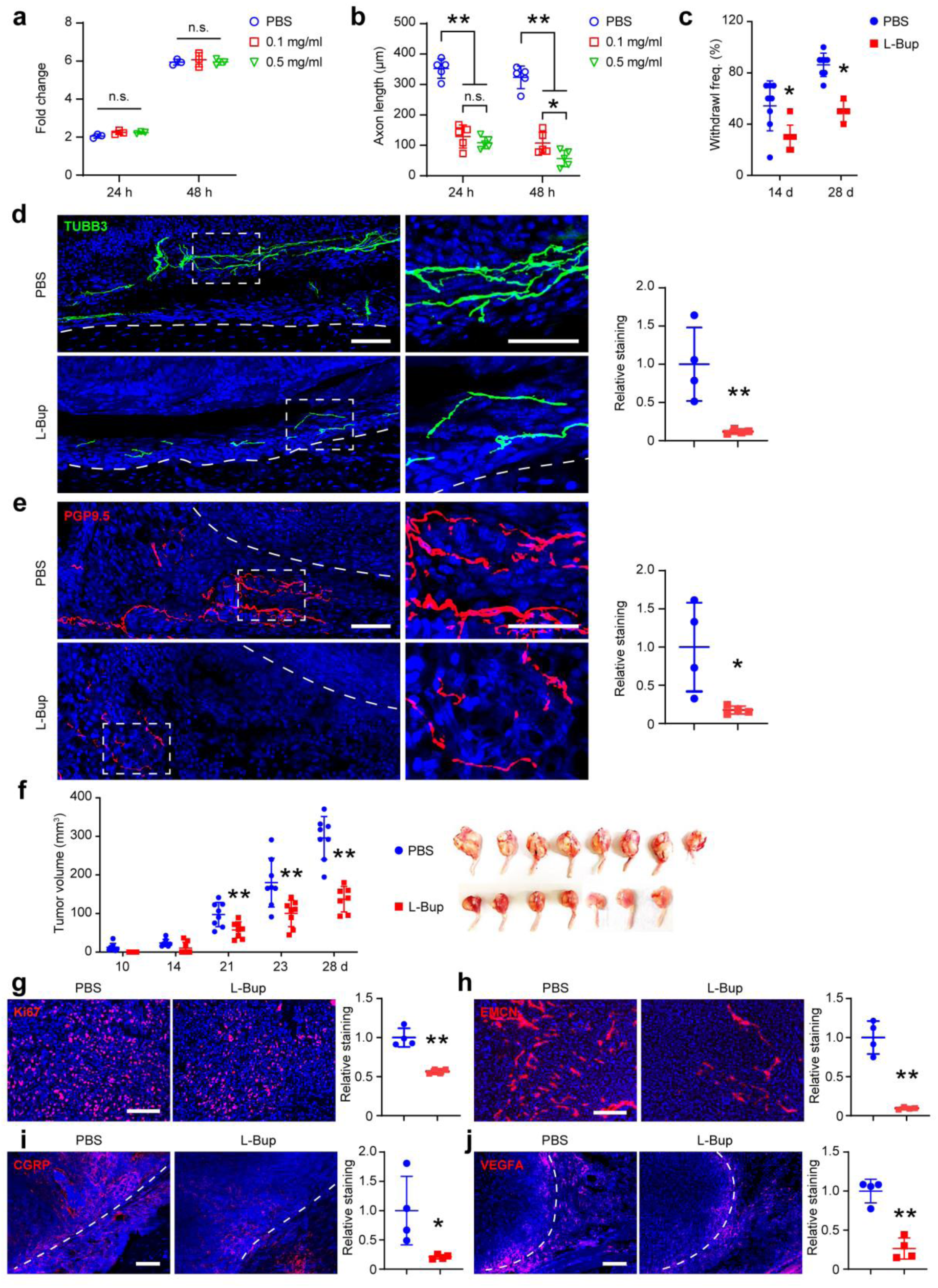
Liposomal Bupivacaine treatment impairs tumors innervation and mitigates OS disease progression. **a**, MTS proliferation assay among 143B OS cells after liposomal bupivacaine (L-Bup) treatment (0.1-0.5 mg/mL, 24 and 48 h). **b**, Axon length measurement among cultured mouse DRG neurons after L-Bup treatment (0.1-0.5 mg/mL, 24 and 48 h). **c-h**, NOD-*Scid* mice were pre-treated with L-Bup or PBS locally and injected with 10^6^ 143B OS cells. All mice were treated with either L-Bup (10 mg/kg) or PBS 2 times / wk for 4 wks. **c**, Von Frey testing of the tumor-bearing limb at 14 and 28 d after tumor implantation among L-Bup or PBS treated mice. **d**, Beta-III Tubulin (TUBB3) immunostaining of peritumoral innervation and quantification among L-Bup or PBS treated mice. **e**, PGP 9.5 immunostaining of sensory neurons and quantification among L-Bup or PBS treated mice. **f**, Tumor volume calculated by caliper measurements twice weekly until 28 d (left) and gross pathology of tumors at 28 d (right). **g**, Ki67 immunostaining and quantification among L-Bup or PBS treated mice. **h**, Endomucin (EMCN) immunostaining and quantification among L-Bup or PBS treated mice. **i**, CGRP immunostaining and quantification among L-Bup or PBS treated mice. **j**, VEGFA immunostaining and quantification among L-Bup or PBS treated mice. Scale bar: 100 µm. Data are shown as the mean ± 1 SD. Individual dots in scatterplots represent values from single animal measurements. N=8 per group in **c**, **f**, and N=4 in **d**, **e**, **g**-**j**. Statistical analysis was performed using unpaired two-way Student’s t test. \**p* < 0.05, \*\**p* < 0.01. Dashed line indicating tumor edge.

## Discussion

The unchecked growth of tumor cells has long been associated in tumors of epithelial origin with disorganized peripheral nerve sprouting^3,39,41–44^. In the present study, we report for the first time that nerve sprouting in sarcoma is functionally relevant, and that these nerves decorate the sarcoma periphery to regulate tumor growth and tumor vascularity. Interestingly, sarcoma tumors were observed to be essentially devoid of nerves within their interior, suggesting a dynamic co-existence of peripheral nerve recruitment and destruction in sarcoma biology.

Neurotrophins and neurotransmitters, such as NGF and CGRP, act directly on tumor cells to promote tumor development. Studies suggest that NGF expression in the tumoral niche is sufficient to recruit nerve fibers in gastric, breast, pancreatic, and oral squamous cell carcinoma. In parallel, nociceptive neurons promote tumor growth by releasing CGRP among other neuropeptides/neurotrophins ^43,45^. Our findings bring up potential therapeutic implications of targeting NGF-TrkA signaling in sarcomas, targeting peripheral neurons via non-specific means, or targeting signaling pathways downstream of neuron-to-sarcoma interactions. Certainly, in epithelial cancers, it has been suggested that targeting NGF has been associated with improved prognosis^46–49^. Although monoclonal antibodies against NGF have been trialed in human patients, a high risk of joint-related adverse events has resulted in stoppage of further clinical development^3,50–53^. Tissue-agnostic approaches such as selective or non-selective inhibition of Trk (larotrectinib and entrectinib) have been approved by the FDA for fusion cancers and could be considered as future therapies to target nerves in sarcoma^54^. More recently, the FDA approval for antibodies against the CGRP (erenumab and fremanezumab) to treat chronic migraines^55–58^, brings up an exciting possibility of drug repurposing to ameliorate sarcoma progression and bone pain. Alternative strategies to perturb nerve-sarcoma interactions are feasible, such as increasing expression of chemorepulsive proteins or other factors^59^. Collectively, abundant potential therapeutic interventions to prevent pathological innervation in sarcomas are feasible and could serve as novel adjunctive therapies in combination with standard chemotherapeutic regimens and surgical resection.

Sensory neurons modulate the tumor microenvironment via secondary mechanisms such as promoting vascularization^60^. In our xenograft orthotopic model of OS, we observed that blocking of sensory (TrkA^+^/CGRP^+^) nerve fibers resulted in marked reduction in tumor-associated blood vessels (EMCN^+^/CD31^+^) and reduced pro-vasculogenic ligand VEGFA, as described in other contexts^18,25,61,62^. These aggregate findings suggest a likelihood that NGF secreting endothelial cells may secondarily contribute to the changes in tumor vasculature as both VEGF-A and NGF sensitize peripheral sensory nerve fibers to several sensory stimuli^63,64^. Although previously best known as a pro-angiogenic molecule, past and recent studies have shown VEGF-A can promote sensory neuronal growth and is neuroprotective^64^. Therefore, such a VEGF-A enriched tumor microenvironment, in combination with other neurosecretory provasculogenic ligands (PDGFA, PDGFB, PGF), may further induce axonal sprouting and survival to facilitate vascular ingrowth directly and indirectly.

Several limitations of this study are worth further investigation and discussion. First, several mouse DRG neuronal subclusters as defined by single cell sequencing approaches are established to express TrkA^65^. Future work may more precisely characterize sarcoma-associated peripheral neurons at the single-cell level and uncover the changes in DRG neuron response over time of sarcoma disease progression. In addition, it is unclear if sensory neurons regulate sarcomas of bone in particular, or if these findings are more broadly relevant to soft tissue sarcomas as well. Lastly, our studies with lipid analgesic nanoparticles were locally applied at the site of tumor growth. It is entirely possible application of drug at the corresponding nerve roots or along the length of the innervating nerve fiber may be a translatable but equally efficacious delivery modality^66^.

Taken together, by applying single-cell technology to orthotopic xenograft implants and their innervating neurons within the dorsal root ganglia, this work has uncovered vital neurosecretory functions of sensory neurons in regulation of OS disease progression. The neural-sarcoma interaction is targetable, and commonly used pharmacotherapies that target peripheral neurons may be leveraged for this purpose. More broadly, multi-tissue sequencing approaches such as this may advance our understanding of neural-cancer interactions, with relevance across diverse malignancies.

## Methods

### Animal use

All animal procedures were conducted in accordance with the guidelines and approved protocol (MO20M128) of the Animal Care and Use Committee (ACUC) at Johns Hopkins University (JHU). NOD-*Scid* mice (Stock No: 001303) were purchased from the Jackson Laboratory (Bar Harbor, ME, USA). TrkA^F592A^ mice were donated from the Ginty laboratory, which are homozygous for a phenylalanine-to-alanine point mutation (F592A) in exon 12 of the mouse *Ntrk1* gene^22^ (Stock No: 022362). This frameshift mutation in TrkA^F592A^ mice renders the endogenous TrkA kinase sensitive to inhibition by the membrane-permeable small-molecule 1NMPP1. Male and female, 8-10 wk-old animals were used for all experiments. *Thy1*-YFP mice, which harbor a transgene derived from the mouse *Thy1* gene that directs expression of YFP in all neurons, are commercially available (Stock No: 003709). For all experiments, TrkA^WT^ and TrkA^F592A^ mice were crossed with *Thy1*-YFP reporter mice. Next, TrkA^WT^;*Thy1*-YFP or TrkA^F592A^;*Thy1*-YFP mice were crossed with NOD-*Scid* mice to allow for xenografting of human tumor cells, used for all experiments (TrkA^WT^;*Thy1*-YFP;NOD-*Scid* and TrkA^F592A^;*Thy1*-YFP;NOD-*Scid)*.

To achieve temporal inhibition of TrkA catalytic activity in TrkA^F592A^ mice, the small molecule 1NMPP1 was used in similarity to prior reports, which was synthesized by Aurora Analytics LLC using standard techniques^11,25^. Purity (99.2%) was confirmed by HPLC-UV254, and characterization by 1H NMR (400 MHz, DMSO-d6) was consistent with structure. Stock solution was prepared at 200 mM by dissolving 1NMPP1 powder in DMSO. For 1NMPP1 administration, i.p. injections were given 48 h before and 2 h before cell implantation using a 5-mM solution at a dosage of 17 μg/g BW. DMSO was used as vehicle control. Animals were thereafter maintained on 1NMPP1-treated drinking water throughout the study period (40 μM 1NMPP1 in ddH2O with 1% PBS–Tween 20). The maximum tumor burden approved by the Ethics Committee was not exceeded in all experiments. The details of the mouse lines used are summarized in **Supplementary Table S13**.

### Osteosarcoma implantation and treatments

Human osteosarcoma cell lines were purchased from American Type Culture Collection (Manassas, VA), 143B (ATCC®-CRL-8303™). Cells were cultured in DMEM supplemented with 15% FBS, 100 U/mL penicillin and 100 μg/mL streptomycin (Gibco, Grand Island, NY) in a humidified incubator with 5% CO2 at 37°C. All animal studies were performed with institutional ACUC approval within Johns Hopkins University, complying with all relevant ethical regulations. For all 143B OS cell implantation, 8-10 wk-old, male and female, TrkA^WT^; NOD-*Scid* or TrkA^F592A^;NOD-*Scid* mice were used. A total of 1 × 10^6^ 143B cells in 50 μl PBS were injected subperiosteally within the left proximal tibia metaphysis under 1.0-1.5% isoflurane in 100% O_2_. Tumor size was measured by caliper twice weekly for four wks, and tumor volume was calculated^23,24^. Primary tumors (*N*=8 mice per group) and lungs (separate cohort, *N*=5 mice per group) were harvested at 28 d post injection for histologic analysis. In a separate cohort of animals, overall survival was evaluated from the first day of tumor cell injection until death using the Kaplan-Meier estimate (*N*=10-13 mice per group). In another cohort of animals, Exparel™ (multivesicular liposomal bupivacaine suspended in 0.9% sodium chloride; 13.3 mg bupivacaine mL^1^), an FDA approved long-acting local analgesic was injected locally in NOD-*Scid* mice. At the same time of tumor inoculation, mice received 10 mg/kg liposomal bupivacaine (L-Bup)^67^ or equal volumes of PBS (*N*=8 mice per group), administered orthotopically every 3 d for four wks using a 25 gauge needle to maintain the structural integrity of the liposomal bupivacaine particles.. Approximately 3% bupivacaine is unbound upon injection, which provides immediate numbness, while the remaining drug is released from the liposomes over time. Pharmacokinetics data suggest that local administration of L-Bup liposomes results in systemic plasma levels of bupivacaine which can persist for 96 hours. Each ml of the drug consists of approximately 37,000,000 multivesicular liposomes/mL. For every 37,000 multivesicular liposomes, approximately 20,000 OS cells were injected.

### Bioluminescence imaging

In a separate cohort of animals, 143B human tumor cells transfected with CMV-Luciferase-RFP-TK Lentivector (BLIV102VA-1) were inoculated for *in vivo* imaging (*N*=5 mice per group)^68^. 28 d post tumor inoculation, mice were anaesthetized with 1.0-1.5% isoflurane, and D-luciferin was injected (150 mg/kg body weight i.p.; 30 mg/ml luciferin) 10 min before imaging. Ventral and dorsal images were collected for 30 s to 2 min using the IVIS imaging system (Xenogen Corp., Alameda, CA). Photons emitted from the primary tumor was quantified using Living Image software (Xenogen Corp.) and total region of interest was calculated with normalized threshold.

### ^18^F-FDG-PET imaging

For ^18^F-fluorodeoxyglucose positron emission tomography (^18^F-FDG-PET), tumor-bearing mice were fasted overnight. Mice were injected with 200 μCi of FDG in 200 μl saline intravenously. 1 h after injection of the radiotracer, a 30-min static whole-body image was acquired. Images were decay corrected and reconstructed using 2-dimensional (2D) OSEM (ordered subset expectation maximization)^69^. After accounting for injected dose and body weight, all radioactivity concentration values were converted into standardized uptake values (SUV). Image analysis was performed using AMIDE software (SourceForge, San Diego, CA) on regions of interest (ROI) drawn over the tumor.

### von Frey testing

Behavioral testing by von Frey filament (0.4 g) stimulus and limb withdrawal was performed at 0, 14, and 28 d after tumor inoculation. Mice were placed in a clear plastic confinement with a custom-manufactured metal mesh platform. Mice were allowed to equilibrate to their surroundings for at least 30 min on the day before and during testing^70^. An indication of the amount of paw withdrawals in the ten trials was expressed as percent response frequency.

### Single cell RNA-sequencing (scRNA-seq)

Xenograft OS tumors were harvested and microdissected 12 d after 143B OS cell inoculation and digested with collagenase type I and II (1 mg/ml; Roche) digestion for 45 min. Cell fractions were collected and resuspended in red blood cell lysis buffer at 37°C for 5 min (Quality Biological, Gaithersburg, MD). Digestions were sequentially filtered through 100-μm and 40-μm sterile strainers. Cells were then washed in PBS and resuspended in Hank’s Balanced Salt Solution (HBSS) at a concentration of 1,000 cells/μl. Cell viability was assessed with Trypan blue exclusion on a Countess II (Thermo Fisher Scientific) automated counter and showed >85% viability across samples. Cells were sent to the JHMI Transcriptomics and Deep Sequencing Core for library preparation. The library was generated using the 10x Genomics Chromium controller following the manufacturer’s protocol^11,18^. Cell suspensions were loaded onto a Chromium Single-Cell A chip along with reverse transcription (RT) master mix and single-cell 3′ gel beads, aiming for 10,000 cells per channel. Following generation of single-cell gel bead in emulsions (GEMs), reverse transcription was performed and the resulting post–GEM-RT product was cleaned up using DynaBeads MyOne Silane beads. The cDNA was amplified, cleaned and quantified using SPRIselect (Beckman Coulter, Brea, CA, USA), and then enzymatically fragmented and size-selected using SPRIselect beads to optimize the cDNA amplicon size before library construction. An additional round of double-sided SPRI bead cleanup was performed after end repair and A-tailing. Indexes were added during PCR amplification, and a final double-sided SPRI cleanup was performed. Libraries were quantified by Kapa qPCR for Illumina Adapters (Roche), and size was determined by Agilent Bioanalyzer 2100. Read 1 primer, read 2 primer, P5, P7, and sample indices (SIs) were incorporated per standard GEM generation and library construction via end repair, A-tailing, adaptor ligation, and PCR. Libraries were generated with unique SIs for each sample. Libraries were sequenced on an Illumina NovaSeq SP 100 cycles (San Diego, CA, USA). CellRanger was used to perform sample demultiplexing, barcode processing, and single-cell gene counting [alignment, barcoding, and unique molecular identifier (UMI) count]. Cells from mouse and human were aligned using two reference genome assemblies mm10 (version M23 Ensembl 98) and GRCh38 (version 32 Ensembl 98), respectively. “CellRanger count” was next used with default settings to align reads to the integrated reference genome and quantify gene expression. Downstream analysis steps were performed using Seurat (v4.4). Cells were first filtered to have >500 and <8,000 detected genes, as well as less than 5% and 25% for mouse mitochondrial and human mitochondrial transcripts, respectively. SCTransform, including regression for cell cycle scores derived using the CellCycleScoring function and dimensional reductions using uniform manifold approximation and projection, was performed. Pathway activation or module scores were generated using the AddModuleScore function of Seurat using validated gene lists from Gene Set Enrichment Analysis (GSEA) and KEGG. Module scores were calculated as the level of gene expression enrichment of a set gene list relative to a random control list, with higher module score values representing positive enrichment control^25,71^. Specific gene lists for module scores are provided in **Supplementary File S1**.

### Histology, immunohistochemistry and image analysis

Primary tumor samples and lungs were fixed in 4% PFA at 4°C for 24 h; tumor samples were processed further for decalcification in 14% EDTA (Sigma-Aldrich, Burlington, Massachusetts) at 4°C under gentle agitation for 28 d. Samples were then soaked in 30% sucrose overnight at 4°C and embedded in optimal cutting temperature compound (OCT, Tissue-Tek 4583, Torrance, CA) for cryosectioning. Routine H&E staining was performed on coronal sections of the lung tissues (15 µm thickness) and number of metastasis foci were manually counted. For immunohistochemistry, sagittal (40 µm thick) sections of primary tumors were washed in PBS × 3 for 10 min and permeabilized with 0.25% Triton-X for 30 min. Next, 3% normal goat serum (Jackson laboratory, Bar Harbor, ME, USA) was applied for 30 min, then incubated in primary antibodies overnight at 4°C (see **Supplementary Table S14)**. The following day, slides were washed in PBS, incubated in the (Vectashield H-1800, Vector Laboratories, Burlingame, CA). Digital images of these sections were captured with 10-40× objectives using inverted fluorescent microscopy (Zeiss LSM 800 and Zeiss LSM 900, Carl Zeiss Microscopy, GmbH). For TUNEL assay, Click-iT™ Plus TUNEL Assay Kits for In Situ Apoptosis Detection kit was used (Thermo Fisher Scientific).

All image analyses were performed using Imaris software 9.3 (Oxford Instruments, Abingdon, England) on maximum projections of 7-15Z-plane sections in a blinded fashion. Semi-automated analyses using Imaris surface generation tool was used to render the surfaces for TUBB3^+^, CGRP^+^, TH^+^, NF200^+^ axons and EMCN^+^ blood vessels. Region of interest (ROI) was set to 500*500 *40 pixels to capture the sarcoma, pseudocapsule (peritumoral area) and the surrounding nerve fibers. To delineate the axon-to-OS spatial relationship, a surface outlining the margins of the sarcoma pseudocapsule was generated based on DAPI staining. A Vantage plot was created to display the relative abstract numerical values reflecting relationships between nerve fibers and tumor cells. Automated quantification of the median distances between individual sarcoma cells to nerve fibers or blood vessels was computed using the Imaris distance transformation tool^72^.

For quantification of aberrant nerve sprouting, three random 20x 3-D volumetric ROI of (300*200 *40 pixels) were analyzed per section per nerve fiber immunostaining, centered around the sarcoma pseudocapsule or periosteum in case of normal bone. A mean value of all three volumetric densities per section is represented as fold increase compared to normal bone.

Digital 3D confocal images of the tumors were captured with a Zeiss LSM800 or Zeis LSM900 microscope (Carl Zeiss) and converted to 2D time-series TIFF stacks by performing z-projections using the inbuilt Zeiss analysis tool or NIH ImageJ software. Then, images were processed by contrast enhancement (stack histogram equalization and normalization, 0.4% saturated pixels). For blood vessel morphometric analysis, images were processed by ImageJ (NIH) and analyzed using the AngioTool (National Cancer Institute) software^73^.

### *In vitro* assessments

143B cell proliferation was assessed using the CellTiter 96 AQueous Nonradioactive MTS Cell Proliferation Assay (Promega). Briefly, 2,000 cells were seeded in 96-well plates and treated with 1NMPP1 (10 μM and 100 μM) or L-Bup (0.1 mg/ml and 0.5 mg/ml) for 24 - 48 h. The optical density at 490 nm was measured on a microplate spectrophotometer (BioTek, Agilent Technologies).

*In vitro* effects of L-Bup (Pacira Pharmaceuticals, Parsippany) on neurite outgrowth were assessed by measuring the axon length. Lumbar DRG neurons (L1-L6) were harvested from 8 wks old female C57BL/6J mice (N=4). Single cell suspension was obtained by digesting DRGs in Collagenase type 1 and Dispase II (1 mg/ml; Roche) at 37°C for 70 min^18^. Cells were filtered at 70 μm and centrifuged at 500 rcf for 5 min. Cells were then mixed with αMEM with 5% FBS, 1 x penicillin/streptomycin, 1 x Glutamax (Thermo Fisher), and anti-mitotic reagents (20 μM 5-fluoro-2-deoxyuridine and 20 μM uridine; Sigma Aldrich). A total of 10,000 cells were seeded into 12-well plate pre-coated with 100 μg/ml poly-D-lysine (Sigma Aldrich) and 10 μg/ml laminin (Thermo Fisher). 48 h later, DRG neuronal cells were treated with L-Bup (0.1 mg/ml and 0.5 mg/ml) for 24 - 48 h. Axon length was measure by ImageJ with NeuronJ plugin^74,75^. Measurements were performed in at least five random fields of three independent repetitions.

### Statistical analysis

All analyses were performed by investigators blinded to the sample identification. Data are presented as mean ± 1 SD. Statistical analyses were performed using GraphPad Prism (version 9.0) or R version 4.1.2. *In vitro* experiments were performed in biologic and experimental triplicate. The number of animals used for *in vivo* experiments is shown in the figure legends. Pilot studies examining tumor volume by caliper measurement among TrkA^F592A^;NOD-*Scid* mice showed large effect sizes of 1.83 or higher. For this scenario, with 8 mice per group a two-sample t-test will provide 80% power to detect effect sizes of at least 1.8 assuming a two-sided 0.05 level of significance. The Kolmogorov-Smirnov test was used to confirm normal distribution of the data. A two-tailed Student’s t test or Wilcoxon test was used for two-group comparisons. A one-way analysis of variance (ANOVA) test was used for multiple groups, followed by Tukey’s multiple comparisons test. \**p* < 0.05, \*\**p* < 0.01, \*\*\**p* < 0.001 and were considered significant.

## Acknowledgements

We thank Dr. Ziyi Wang and the JHU microscopy facility and JHMI Transcriptomics and Deep Sequencing core for their technical assistance. Funding to AWJ includes NIH/NIAMS (P01 AG066603, R01 AR079171, R21 AR078919), NIH/NIDCR (R01 DE031488, R01 DE031028), Alex’s Lemonade Stand Foundation (22-26743), American Cancer Society (DBG-23-1155131-01-IBCD), the Maryland Stem Cell Research Foundation (2021-MSCRFD-5641), and Department of Defense (USAMRAA HT9425-24-1-0051). Funding to YG includes NIH/NINDS (NS110598, NS117761). Funding to TJP includes NIH grants (U19 NS130608 and R01 NS 111929). The content is solely the responsibility of the authors and does not necessarily represent the official views of the National Institute of Health, Department of Defense, or U.S. Army.

## Author contributions

Q.Q., S.R., and A.W.J. designed the study; Q.Q., and S.R. performed the experiments, analyzed the data, and were responsible for writing the original draft; Z.L., L.Z., M.C., M.A., X.X., N.T., M.G.S., M.X., M.Z., and L.C. performed the experiments; A.U., and Y.G. developed the methodology for behavioral test. K.M., and T.P. provided single-cell transcriptomics and RNA sequencing data of human DRG neurons. S.R., Z.L., K.M., M.M. and B.L. performed computational analyses. C.D.M and E.F.M. provided human OS biopsy samples. A.W.J and T.L.C. provided and assisted with transgenic mice lines. A.W.J. secured funding, supervised the overall study, handled review and editing of the manuscript. All authors reviewed, commented and approved the manuscript.

## Data availability

Requests for data and materials should be addressed to A.W.J. Single-cell RNA sequencing data of mouse xenograft OS implants generated from this study are deposited in NCBI Gene Expression Omnibus (accession # GSE252589). Human OS cell line data was generated using Cancer Cell Line Encyclopedia RNA sequencing EMBL-EBI database. scRNA-seq of human primary OS dataset was obtained from GEO accession #GSE162454. Human sNuc-seq DRG datasets were obtained from GEO accession #GSE168243 and GSE255436. Human DRG spatial sequencing dataset was obtained from dbGaP repository (phs001158), and the Loupe Browser files are available at <u>sensoryomics.com</u>. Bulk RNA-seq of human OS bones and human DRG neurons with neuropathic-like cancer pain were obtained from GEO (accession # GSE99671) and dbGaP repository (phs001158.v2.p1), respectively. All other relevant data supporting the findings of this study are included within the article. Source data are provided.

## Extended Data Figures

**Extended Data Fig. 1.**
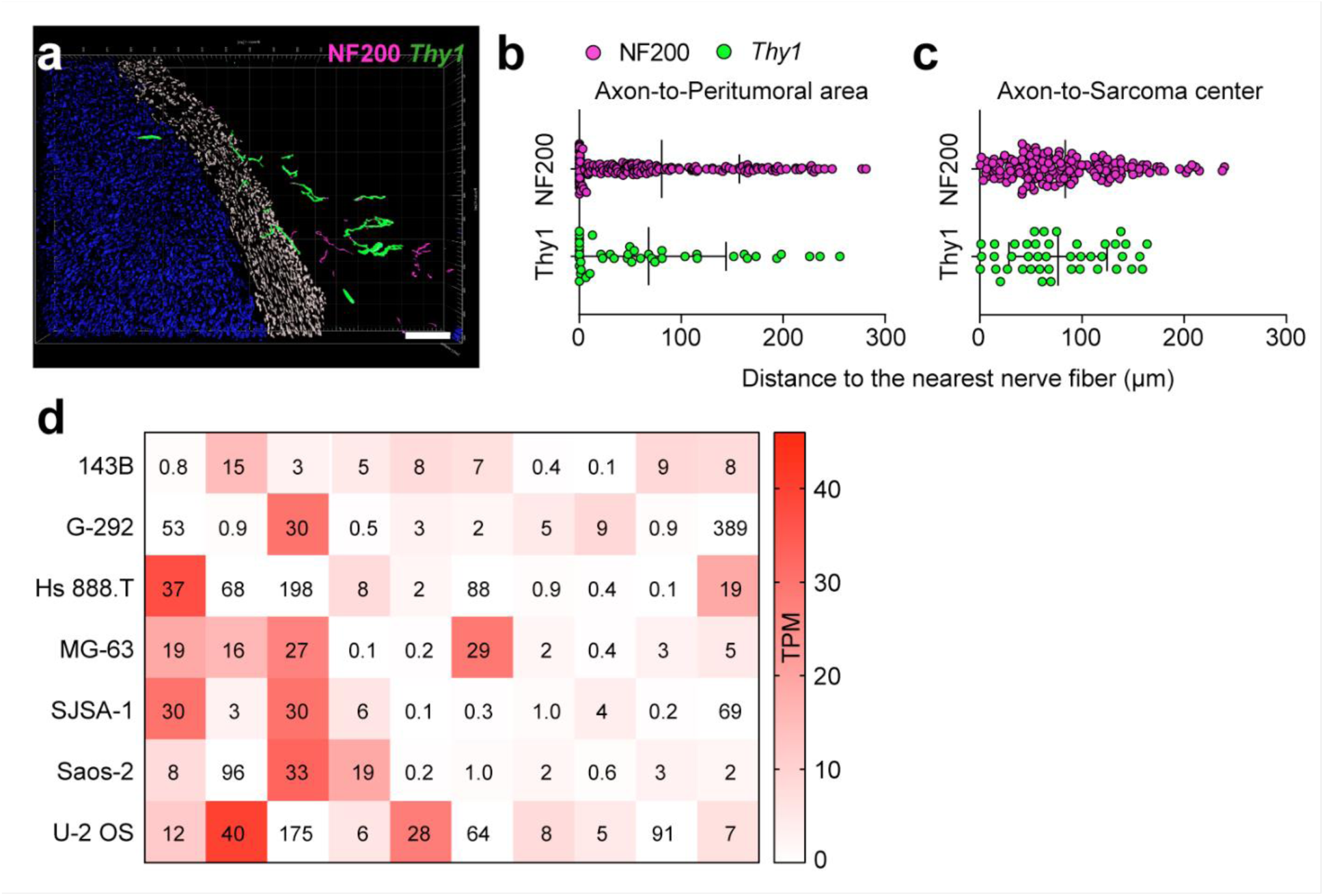
Spatial mapping of nerve-sarcoma interaction (Fig. 1h) and axon guidance molecule expression in tumor cell lines (Fig.1k). **a**, Representative vantage plot of NF200^+^ (magenta) and *Thy1*+ (green) nerve fibers surrounding osteosarcoma (blue) and the sarcoma pseudocapsule (white). **b**, **c,** Scatter plot represents the quantification of the median distance to the nerve fibers from the **b** nearest peritumoral area and **c** sarcoma center. Each dot represents the nearest distance to a single nerve fiber (NF200^+^: magenta, *Thy1*^+^: green). Dots on the zero mark indicates the presence of the nerve fibers on the peritumoral or sarcoma center. Error bar represents median ± 95% C.I. N=3 images used for quantification. Scale bar: 100 µm. **d**, Expression of axon guidance molecules within human osteosarcoma cell lines. Data obtained from EMBL-EBI database^19,20^.

**Extended Data Fig. 2.**
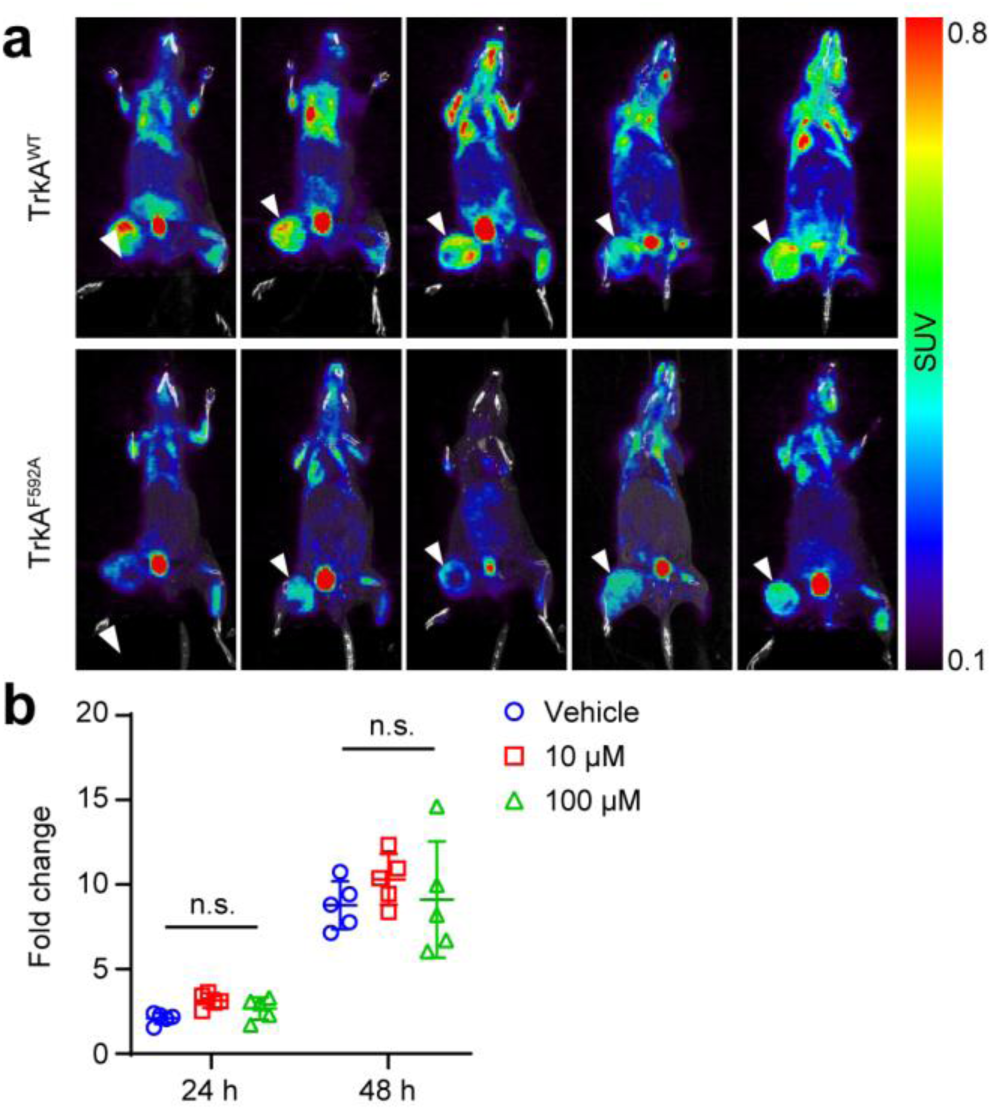
PET-CT images of individual animals (Fig. 2d) and small molecule 1NMPP1 has no direct effects on 143B cell growth *in vitro*. **a,** 143B osteosarcoma implants among TrkA^WT^ or TrkA^F592A^ mice, imaged at 28 d post-inoculation. White arrowheads show the tumor. N=5 mice per group. **b,** 143B cells were treated with 1NMPP1 at indicated dose for 24 and 48 h. Cell proliferation was assayed by MTS assay. Data shown as mean ± 1 SD, indicated by crosshairs and whiskers. Individual dots in scatterplots represent values from single measurements. One way ANOVA followed by Tukey’s test was used for comparison. n.s.: not significant.

**Extended Data Fig. 3.**
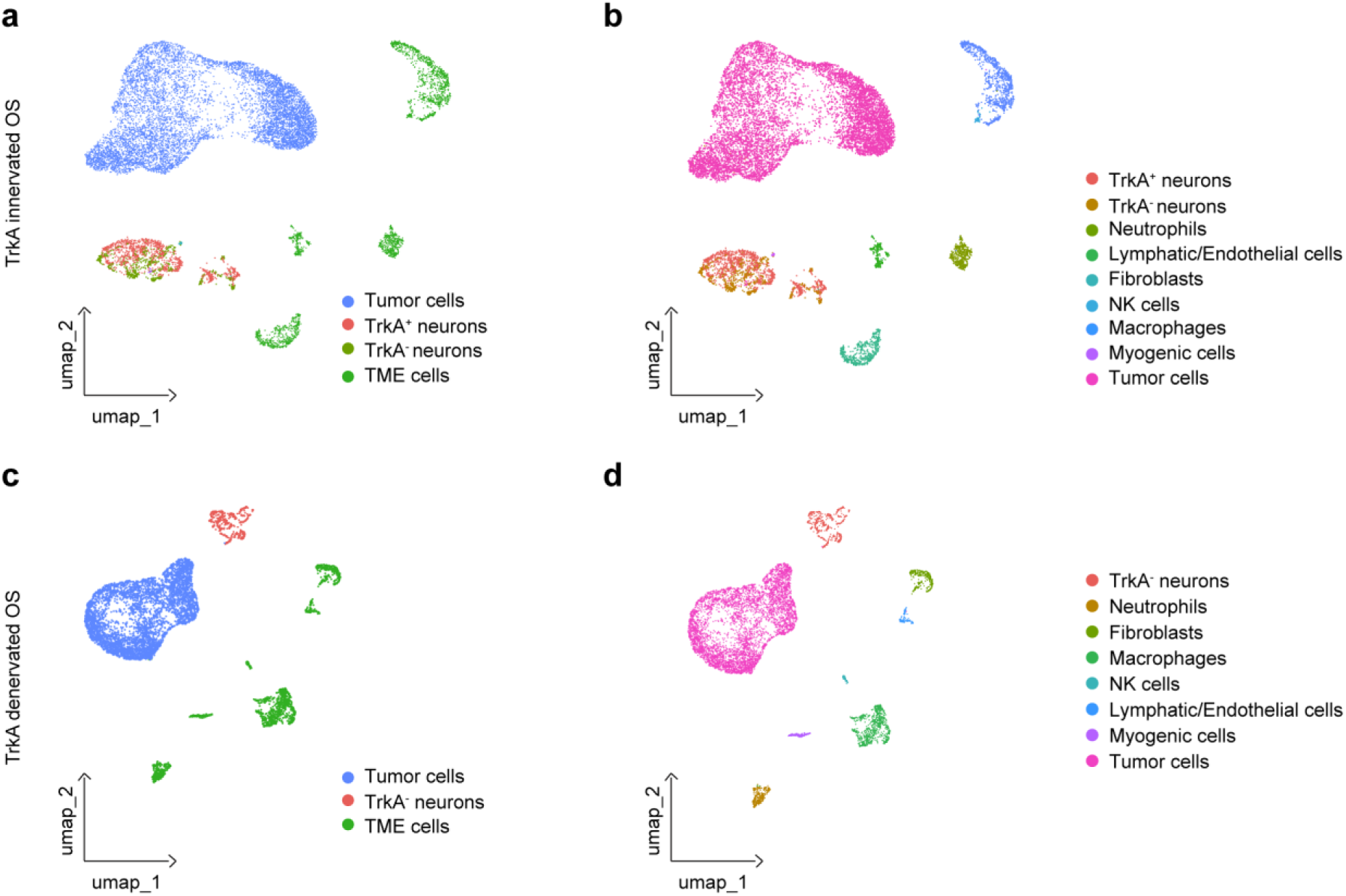
Computational modeling of TrkA innervated and denervated OS niche (Fig. 3h, i). **a,** UMAP and clustering of human lumbar DRG neurons^26^ (TrkA^+^ neurons) with tumor cells and mouse tumor-associated cells identified by **b,** cell type from TrkA^WT^ implants. **c,** UMAP and clustering of human lumbar DRG neurons (TrkA^-^ neurons) with tumor cells and mouse tumor-associated cells identified by **d,** cell type from TrkA^F592A^ implants. N=8,344 Tumor cells, N=2,035 mouse tumor niche cells, N=1,155 TrkA^+^ neurons, and N=602 TrkA^-^ neurons.

**Extended Data Fig. 4.**
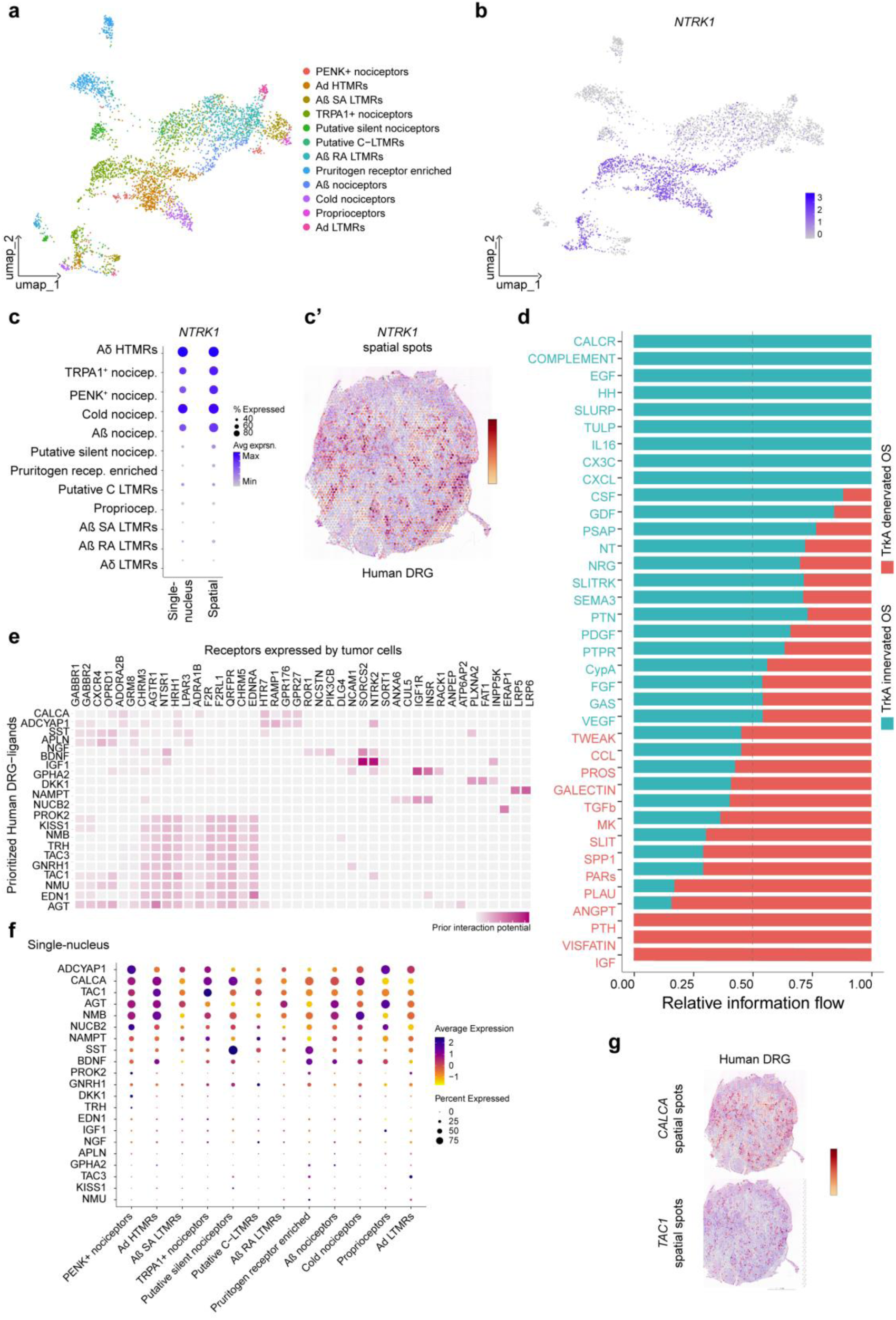
Overview of single-nucleus RNA sequencing and spatial transcriptomics of human DRG neurons (Fig. 3). **a,** UMAP and clustering of human DRG neurons^28^. **b,** Feature plot of *NTRK1*. Dot plot of *NTRK1* by **c,** single nucleus^28^ and spatial^29^ sequencing. **c’,** Spatial localization of *NTRK1*. **d,** Relative information flow among TrkA innervated and TrkA denervated OS. **e,** NicheNet analysis of human DRG^28^ derived ligands with gene targets among tumor cells derived from TrkA^WT^ implants. **f,** Dot plot showing expression of top-ranked DRG ligands among various human DRG neuronal^28^ subclusters. **g,** Spatial localization ^29^ of *CALCA* and *TAC1* expression among representative human DRG tissue section.

**Extended Data Fig. 5.**
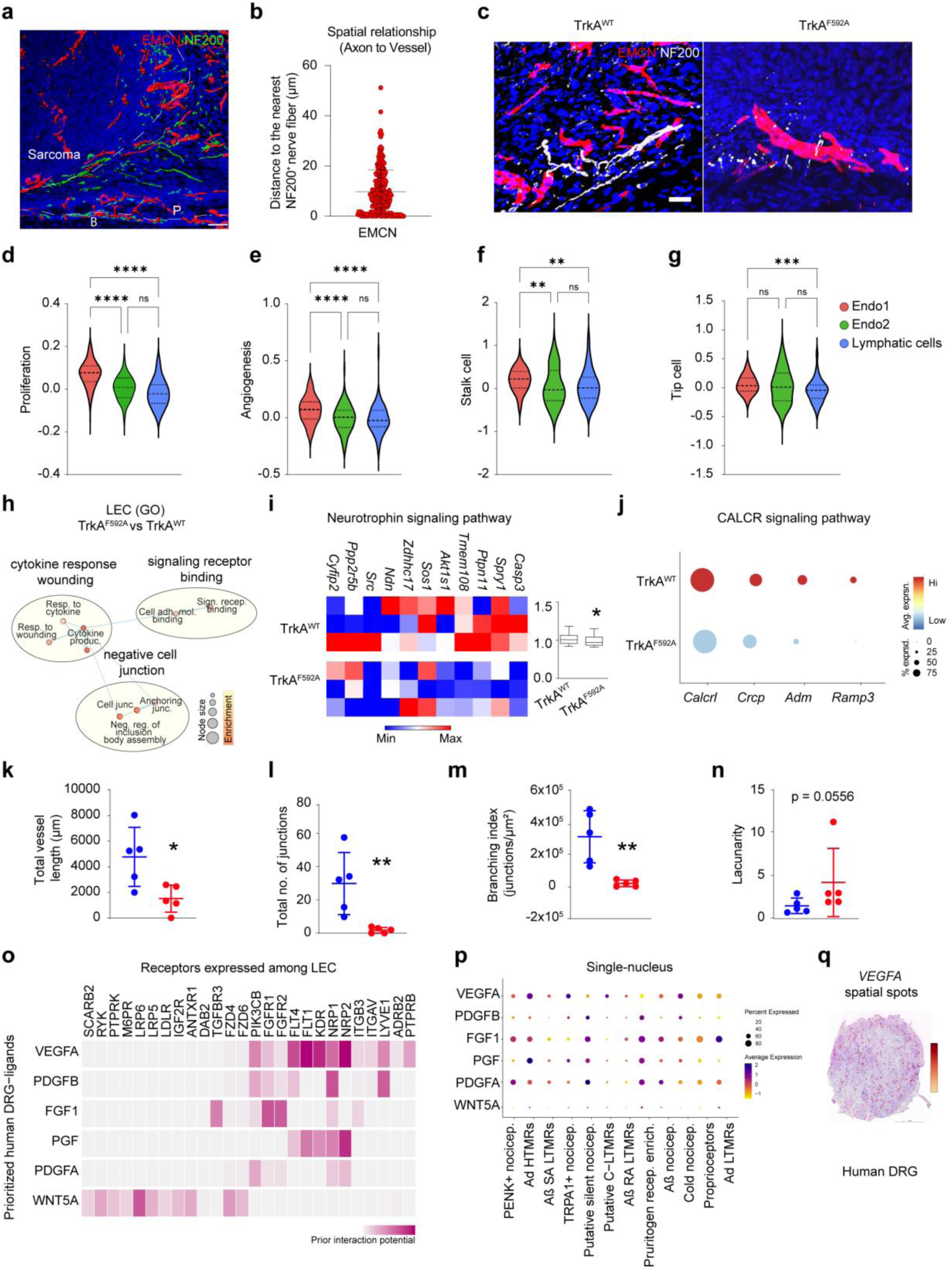
Additional characterization of neurovascular perturbations among TrkA^WT^ and TrkA^F592A^ sarcoma implants (Fig. 4). **a,** Imaris surface renderings of immunohistochemical staining to illustrate the interaction between nerve (NF200^+^, green) and blood vessels (EMCN^+^, red) in a Nod-*Scid* xenograft OS model. The thick dashed white line represents the tumor boundary (OS), the boundary between the periosteum (P) and the underlying cortical bone (B). **b,** Scatter plot represents the quantification of the nearest distance (median) to the nerve fibers from the blood vessels. Each dot represents the nearest distance to a single nerve fiber (NF200^+^, red dot). **c,** Dual immunostaining for EMCN (red) and NF200 (white) among TrkA^WT^ and TrkA^F592A^ tumor implants. Violin plot of module scores among lymphatic/endothelial subclusters, including **d,** cellular proliferation, **e,** angiogenesis, **f,** stalk-cell markers, and **g,** tip-cell markers. Gene lists for module scores are shown in Supplementary File S1. **h,** Network of enriched GO terms and Reactome pathways generated with g:Profiler and EnrichmentMap in Cytoscape among LEC cells from TrkA^F592A^ implants. Each node (circle) represents a gene set characterized by a particular GO term or reactome pathway. Node fill indicates the enrichment score (FDR q-value). The thickness of blue lines (edges) indicates the number of shared genes (overlap) between two connected nodes. Nodes with high overlap are clustered together, forming groups characterized by similar terms and pathways. **i,** Heatmap and modular index scoring of neurotrophin signaling pathway among TrkA^WT^ and TrkA^F592A^ LECs. **j,** CALCR signaling genes among TrkA^WT^ and TrkA^F592A^ LEC, as shown by dot plot. **k-n** Vascular histomorphometric analysis based on CD31 immunostaining, including **k,** total vessel length, **l,** total number of junctions, m, branching index, and **n,** lacunarity among TrkA^WT^ and TrkA^F592A^ tumor implants. **o,** Interactome analysis showing predicted interaction potential among various neural^28^ derived vasculogenic ligands with expressed receptors on TrkA^WT^ LECs. **p,** Dot plot showing expression of top-ranked DRG derived vasculogenic ligands among various human DRG neuronal subclusters^28^. **q,** Spatial distribution^29^ of *VEGFA* among representative human DRG tissue sections. White scale bar: 100 µm in **a**; white scale bar in part **c**, 50 µm. For all graphs, data are expressed as the mean ± 1 SD. \**p* < 0.05, \*\**p* < 0.01, \*\*\**p* < 0.001, \*\*\*\**p* < 0.0001.

**Extended Data Fig. 6.**
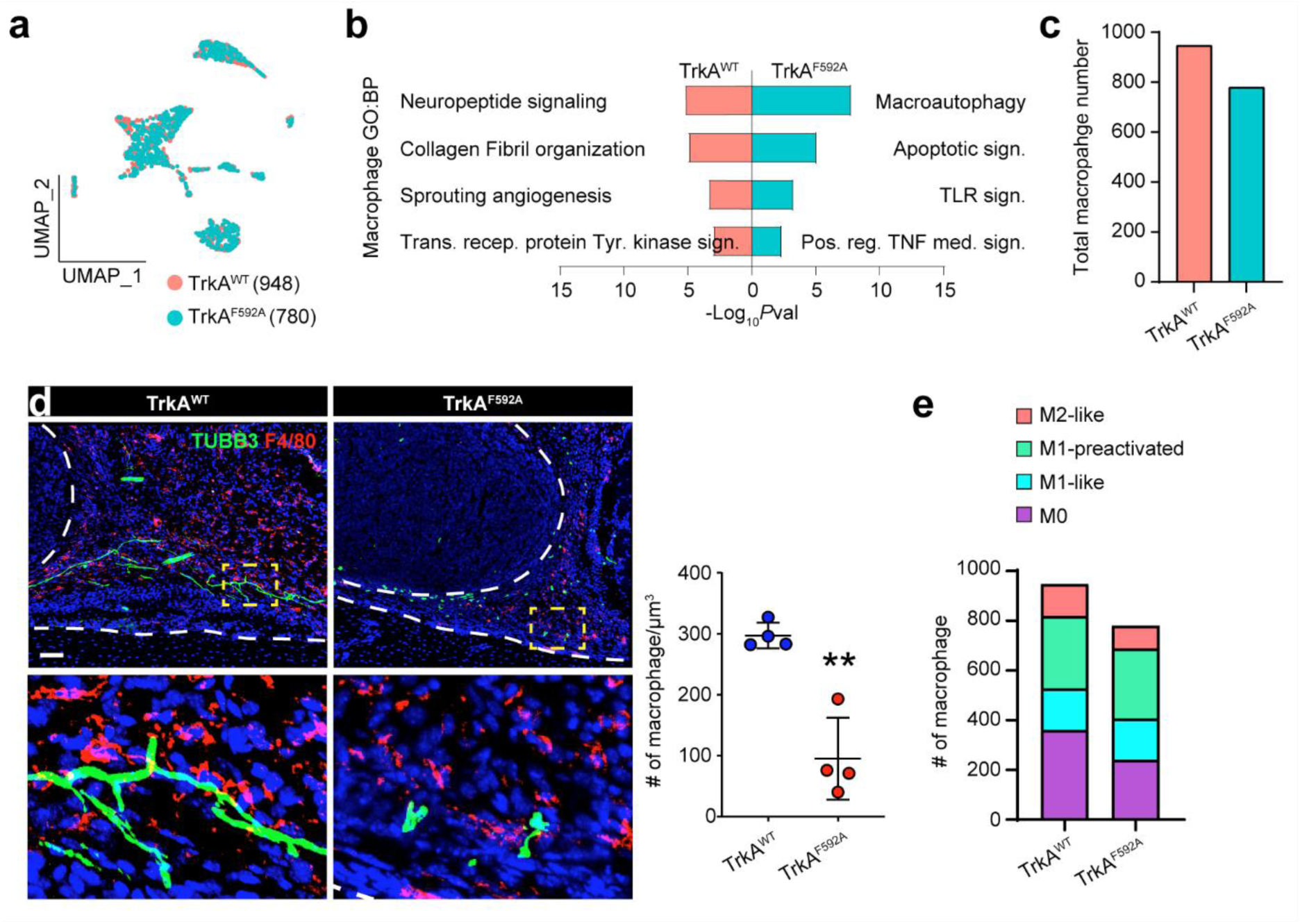
TrkA inhibition impedes sarcoma infiltration by macrophages. **a,** UMAP visualization of macrophage clusters among TrkA^WT^ and TrkA^F592A^ animals. **b,** Gene Ontology (GO) term analyses of significantly enriched biological processes in macrophages among TrkA^WT^ or TrkA^F592A^ mice. **c,** Overall frequency of macrophages by sc-RNA Seq among TrkA^WT^ or TrkA^F592A^ mice. **d,** Macrophage infiltration by F4/80 immunostaining and semi-quantification. **e,** Frequency of macrophage subpopulation stratified by polarization states among TrkA^WT^ or TrkA^F592A^ mice (defined by MacSpectrum software). Scale bar: 100 μm. N=5. Data are expressed as the mean ± 1 SD. Individual dots in scatterplots represent values from single animal measurements. Statistical analysis was performed using unpaired two-way Student’s t test. \*\**p* < 0.01.

## Supplementary Figures

**Supplementary Fig. S1.**
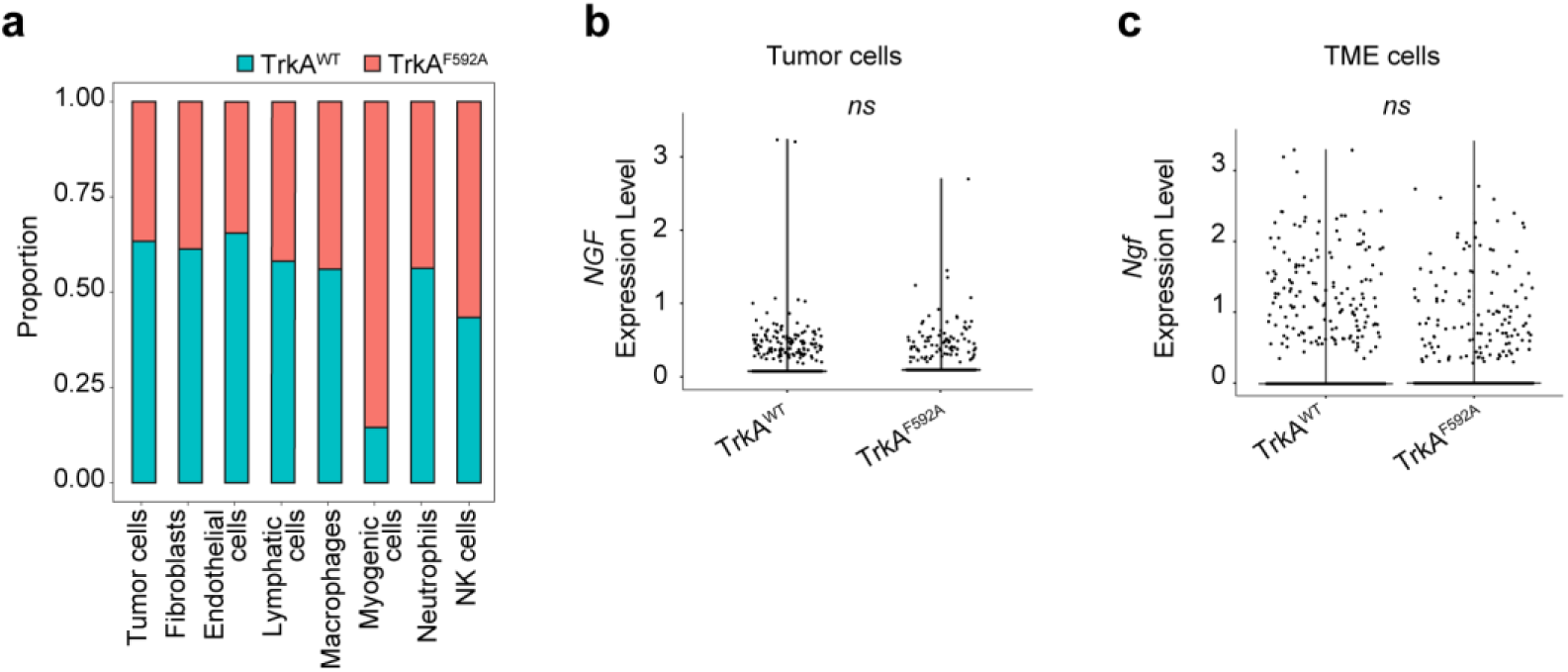
Cell cluster proportion, and Nerve growth factor (*Ngf*) expression within the tumor cells and TME cells harvested from TrkA^WT^ and TrkA^F592A^ by scRNA-seq. **a,** Proportion of cells among each cluster obtained either from TrkA^WT^ or TrkA^F592A^ mice. A value of 0.50 indicates equal cell numbers between TrkA^WT^ and TrkA^F592A^ mice. Violin plot of *Ngf* expression among all **b,** tumor cells and **c,** TME cells. Cells isolated from n=3 mice per genotype.

**Supplementary Fig. S2.**
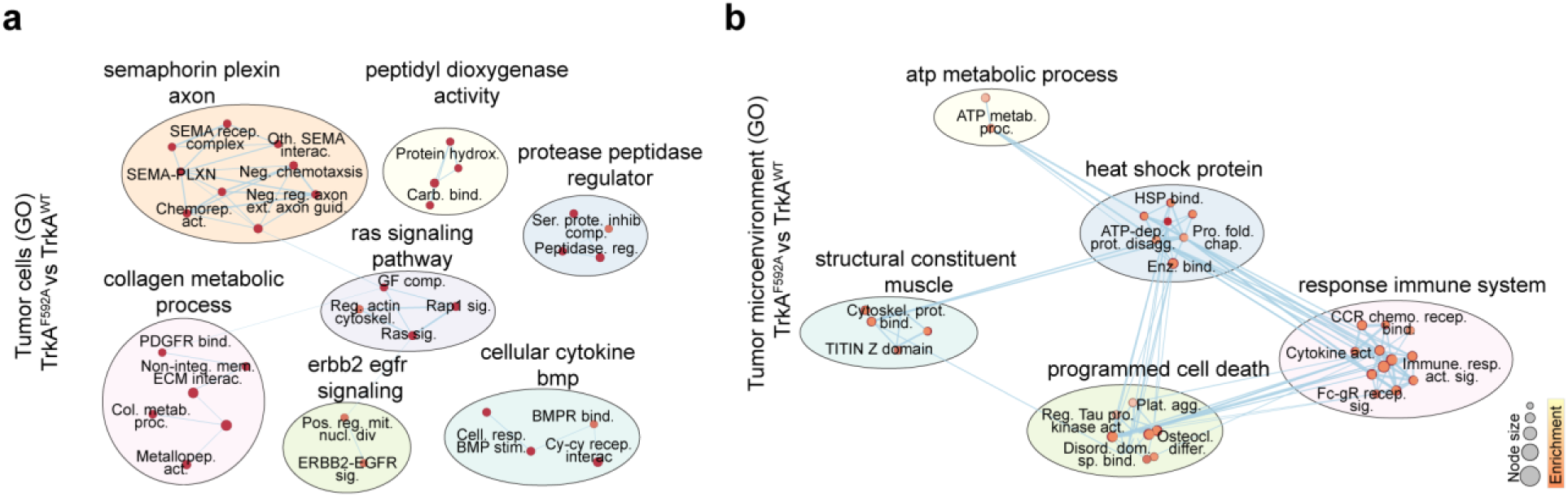
Network of enriched GO terms and reactome pathways generated with g:Profiler and EnrichmentMap in Cytoscape among TrkA^F592A^ implants. **a,** Tumor cells implanted in TrkA^F592A^ mice, and **b,** TME cells from TrkA^F592A^ mice in comparison to TrkA^WT^ mice. Each node (circle) represents a gene set characterized by a particular GO term or reactome pathway. The node fill indicates the enrichment score (FDR q-value). The thickness of blue lines (edges) indicates the number of shared genes (overlap) between two connected nodes. Nodes with high overlap are clustered together, forming groups characterized by similar terms and pathways.

**Supplementary Fig. S3.**
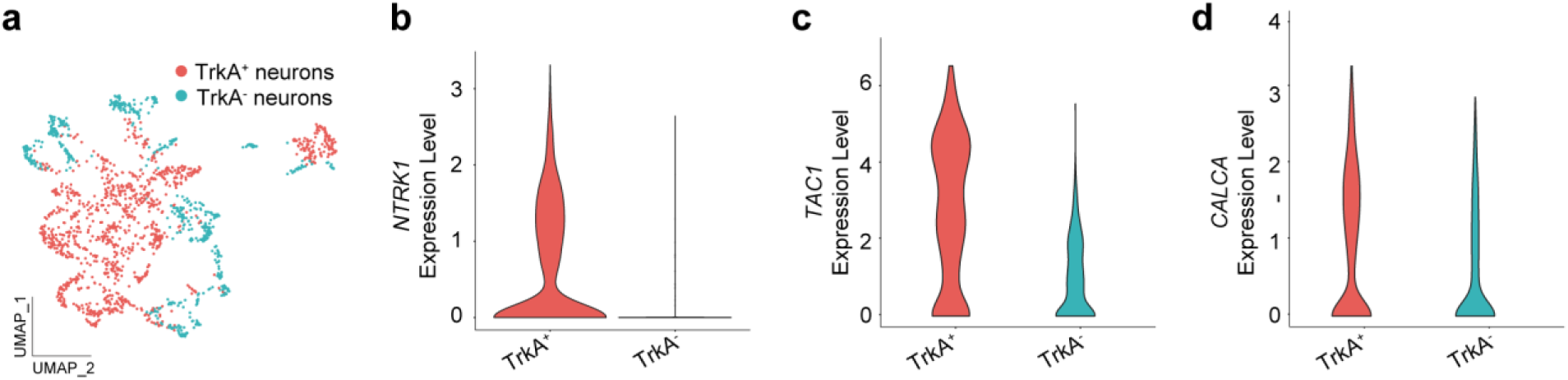
Overview of single-nucleus RNA sequencing (sNuc-seq) of lumbar dorsal root ganglion (Fig. 3h, i). **a,** UMAP plot of DRG neurons^26^ stratified by TrkA expression. Violin plots of **b,** *NTRK1*, **c,** *TAC1*, **d,** *CALCA* among TrkA^+^ and TrkA^-^ neuronal subclusters.

**Supplementary Fig. S4.**
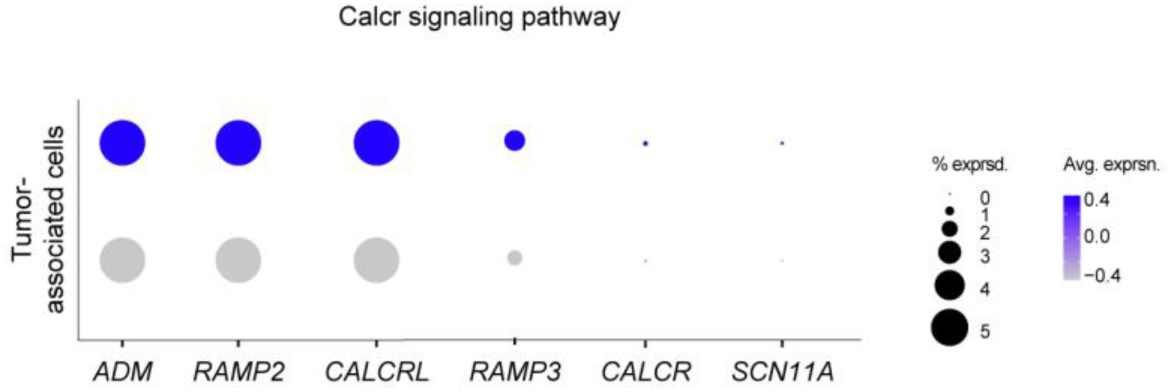
Dot plot of individual CALCR signaling genes among TrkA^WT^ and TrkA^F592A^ TME cells. Single cell RNA sequencing of OS tumor site obtained 12 days after transplantation of 143B OS tumor cells. N=2,242 and 1,802 TME cells analyzed among TrkA^WT^ and TrkA^F952A^ mice, respectively. N=3 mice per group.

**Supplementary Fig. S5.**
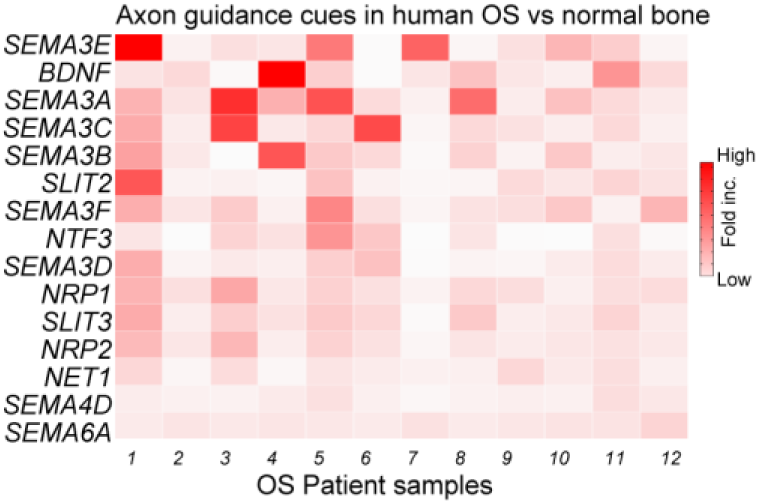
Heatmap of additional axon guidance cues within clinical human OS samples^33^. (*N*=12) compared to normal human bone.

